# Seeing Just Enough: The Contribution of Hands, Objects and Visual Features to Egocentric Action Recognition

**DOI:** 10.64898/2026.02.15.705896

**Authors:** Filip Rybansky, Sadegh Rahmaniboldaji, Andrew Gilbert, Frank Guerin, Anya C. Hurlbert, Quoc C. Vuong

**Affiliations:** Newcastle University; University of Surrey

**Keywords:** Action recognition, egocentric action, minimal video, hierarchical feature extraction, spatiotemporal features, SHAP, feature importance, semantic modelling

## Abstract

Humans recognize everyday actions without conscious effort despite challenges such as poor viewing conditions and visual similarity between actions. Yet the visual features contributing to action recognition remain unclear. To address this, we combined semantic modelling and feature reduction methods to identify critical features for recognizing actions from challenging egocentric perspectives. We first identified egocentric action videos from home environments that a motion-focused action classification network could correctly classify (*Easy* videos) or not (*Hard* videos). In Experiment 1, participants (*N*=136) labelled the action and object in the videos. Using a language model framework, we derived human ground truth labels for each video and quantified its recognition consistency based on semantic similarity. Participants recognized actions and objects in *Easy* videos more consistently than in *Hard* videos. In Experiment 2, we recursively reduced the *Easy* and *Hard* videos with high recognition consistency to extract minimal recognizable configurations (MIRCs), in which any further spatial or temporal reductions disrupted recognition. The data was collected using a large-scale online study (*N*=4360). We extracted information related to the hand, objects, scene background and visual features (e.g., orientation or motion signals) from the 474 MIRCs. Binary classification showed that recognition was disrupted when regions containing the manipulated object and strong orientation signals were removed, while temporal reduction by frame-scrambling disrupted recognition in 73% of MIRCs. The active hand had some marginal contribution. Our results highlight the importance of both mid- and high-level information for egocentric action recognition and link hierarchical feature theories with naturalistic human perception.

## 1. Introduction

Humans quickly and accurately recognize actions in everyday life. This is notable because actions may be seen under poor conditions (e.g., low lighting). Actions can also be visually similar, yet their underlying purpose and outcome can be very different (e.g., hugging vs fighting). Despite advances in computer vision, humans can outperform state-of-the-art action classification systems, particularly for visually similar actions or when there is poor environmental lighting, occlusions or clutter.

Our primary aim in the present study was to determine the critical spatial and spatiotemporal features for action recognition from a challenging egocentric perspective. Previous studies focused on allocentric, third-person actions and it remains unclear whether their findings generalize to egocentric actions. Although allocentric and egocentric views of actions have similarities, there are also important differences in the spatial and spatiotemporal information that is available. Here we developed a novel language-model approach to assess human accuracy at freely naming a wide range of typical home actions presented from an egocentric perspective. We then combine this approach with a feature reduction paradigm to identify minimal recognizable configurations that contain candidate mid- and high-level features for egocentric action recognition.

### 1.1 Critical Features for Action Recognition

What are candidate critical features that are informative or diagnostic for action recognition? A prominent view of visual processing proposes that visual recognition occurs through hierarchical feature extraction, in which visual features are extracted from the scene and mapped onto representations of objects and actions stored in memory. The hierarchical feature extraction employs a complex combination of feed-forward and feed-back processes (Ahissar & Hochstein, 2004; Bar, 2003; Hochstein & Ahissar, 2002), handling spatial, temporal and spatiotemporal information (Ben-Yosef et al., 2020; Casile & Giese, 2005; Giese & Poggio, 2003; Smekal et al., 2024).

Within the hierarchical framework, low-, mid- and high-level features can provide diagnostic information for action recognition. For low- to mid-level features, studies have mainly focused on global and localized temporal motion features encoded in the optic flow of biological motion stimuli (Casile & Giese, 2005; Giese & Poggio, 2003; Hirai & Senju, 2020; Thornton & Vuong, 2004; Troje & Westhoff, 2006). For high-level visual features, on the other hand, researchers suggested sequences of form snapshots ‘integrated across 100ms or more’ (Casile & Giese, 2005; Platonov & Orban, 2016).

Orban et al. (2021) further suggested two classes of high-level spatiotemporal features that can contribute to the recognition of manipulative actions. One feature type corresponds to movements of body parts, and the other to their effects on objects in the environment. For manipulative actions involving hand-object interactions (e.g., washing dishes), empirical evidence suggests that people are sensitive to local features associated with the hand or objects (Loucks & Baldwin, 2009; Flanagan & Johansson, 2003). That is, the critical featural information differentiates the smallest units of action, such as grasp, place, push, and pull, mainly via the hand shape and hand-object relationship or contact (Ben-Yosef & Ullman, 2018; Loucks & Baldwin, 2009).

### 1.2 Egocentric Actions

Egocentric actions are, by definition, viewed from the perspective of the person performing the action. The spatial and spatiotemporal features in egocentric action videos share some similarities with those in allocentric videos (Frischen et al., 2009), but differ in important ways, not least because the camera’s viewpoint is attached to the moving actor. Thus, despite progress in identifying the critical features for action recognition in allocentric videos (e.g., Ben-Yosef et al., 2020; Loucks & Baldwin, 2009; Smekal et al., 2024), it is unclear to what extent findings generalize to egocentric actions. Egocentric action recognition remains a challenging problem for computer vision, and solutions have important applications in autonomous robotics, as well as context-aware interaction with AR/VR and AI assisted healthcare and skill learning.

Here we focused on egocentric actions that involve hand-object interactions. Compared with allocentric views, egocentric perspectives of these actions have greater variability in the visual input, motion, blur and clutter, due to the actor’s moving viewpoint (Ren & Gu, 2010; Swallow et al., 2018). Yet other factors may provide advantages for recognition and strengthen the sensitivity for hand-object interaction: objects within the actor’s affordance are viewed from a closer distance, and their physical properties, identity, potential manipulations and touch locations are more easily accessible (Borghi et al., 2012; Roche & Chainay, 2013). Egocentric perspective also contains more information about grip and may benefit more from temporal information (Campanella et al., 2011). For humans, egocentric actions may elicit increased activation of motor and premotor areas and mirror neurons (Caggiano et al., 2009), enhancing recognition through its simulation matching component (Rizzolatti & Craighero, 2004).

Ben-Yosef et al. (2020) recently used a reduction procedure that recursively reduced the frame size, resolution or number of frames until the original action video from an allocentric perspective could no longer be reliably recognized (see also Ullman et al., 2016 for static images). They found a level at which further reduction of any type would lead to a recognition breakdown; that is, participants could no longer recognize the action in any of the reduced stimuli. In fact, there would often be a large recognition gap between human performance at this level compared to the next level. We refer to the reduced video at the level before the recognition gap as the Minimal Recognizable Configuration (MIRC), corresponding to Ben-Yosef’s minimal video. The recognition gap suggests that MIRCs are densely packed with candidate critical features that can be used to recognize actions (see also Ben-Yosef et al., 2018). Ben-Yosef further showed that computer-vision models had smaller recognition gaps than humans, suggesting that particularly spatial reductions had a more gradual effect on allocentric action recognition for these models.

We recently adapted Ben-Yosef et al.’s (2020) reduction procedure to identify MIRCs for egocentric videos of actions from a home environment in the Epic-ReduAct dataset (Rahmaniboldaji et al., 2025). We also compared recognition parameters between humans and computer-network models. Similar to Ben-Yosef and colleagues (see also Ullman et al., 2016), we found that humans demonstrated a sudden large recognition gap at later reduction levels whereas our computer-vision model demonstrated a gradual decrease and even increase in confidence for egocentric action recognition (Rahmaniboldaji et al., 2025). Here, we will build on this previous work by comparing recognition parameters between *Easy* and *Hard* videos, incorporating a temporal manipulation, and identifying the critical spatiotemporal features for egocentric action recognition in human observers.

### 1.3 Aim and Hypotheses

The primary aim in the current study was to determine critical spatial and spatiotemporal features to recognize actions carried out in a home environment from a challenging egocentric perspective. For this aim, we adapted Ben-Yosef et al.’s (2020) recursive reduction procedure for egocentric action videos selected from the large-scale Epic-Kitchens-100 dataset (EK100; Damen et al., 2022). We divided videos selected for our dataset into *Easy* and *Hard* videos, according to whether they were classified accurately or inaccurately, respectively, by our Motion-Focused Self-Supervised network (MOFO; Ahmadian et al., 2023; see also Rahmaniboldaji et al., 2025). This allowed us to present participants with a wide range of actions performed in real-world home environments, with different variations of similar actions, and with some actions that could not be recognized by abstract features based on MOFO.

In Experiment 1, we used a language-model (LM) approach to analyze individual responses to determine Human Ground Truth (HGT) responses for each action based on the entire video (see also Smekal et al., 2024). In Experiment 2, we implemented the recursive reduction procedure. We measured semantic similarity between participants’ free-naming responses and the HGT labels from Experiment 1, to determine recognition accuracy and establish recognition thresholds defining MIRCs. Lastly, we used tree-based classifiers to determine mid- to high-level spatial and spatiotemporal features from these MIRCs that can be used to recognize action videos from a challenging egocentric perspective, and assess whether these features can help explain why computer-vision networks failed to recognize some videos (i.e., *Hard* videos).

Based on the work on features of allocentric action recognition (e.g., Loucks & Baldwin, 2009), we hypothesized that the hands and objects directly involved in the action would be more important for their recognition than other high-level objects present in the scene. Motion information has been shown to be important for recognition of other people’s actions (e.g., Thornton et al., 1998), so we further expected to find that motion would be more important than other mid-level features. Related to motion cues, we hypothesized that scrambling the order of frames of MIRCs would lead to a recognition breakdown, and more so in MIRCs where the hands and motion were more visible.

## 2. Experiment 1

The purpose of Experiment 1 was to determine a robust HGT label for a large subset of the EK100 dataset (Damen et al., 2022). This also allowed us to compare human performance and a top-performing computer-vision model (MOFO; Ahmadian et al., 2023) in recognizing egocentric actions, and assess accuracy for recognizing MIRCs in Experiment 2.

Our EK-HGT dataset consisted of *Easy* and *Hard* videos selected from EK100 (see below). To determine HGT for these videos, we conducted an online behavioral experiment in which participants typed brief descriptions of the action. We then constructed an LM pipeline to analyze semantic distances between individual human responses and derive robust HGT for each video (Mikolov et al., 2013; Reimers & Gurevych, 2019; Smekal et al., 2024). The HGT labels underpin a metric for (a) quantifying response consistency and accuracy of depicted actions across participants; and (b) determining alignment of human response consistency with classification performance of MOFO as proxied by the *Easy* and *Hard* videos.

### 2.1 Construction of *Easy* and *Hard* video datasets

Videos in the EK100 dataset were recorded from an egocentric perspective and consist of natural interactions with objects in the kitchen (Damen et al., 2022). Actors wore a head-mounted camera and performed daily home activities (e.g., ‘washing dishes’; ‘pouring milk’) so that only the hands, arms and local scene (e.g., nearby objects) were visible. The original EK100 dataset contained human ground-truth labels obtained through transcriptions of actor narrations derived by consensus of three Amazon Mechanical Turk transcribers, which we refer to as the EK100-GT labels. This is a small number of transcribers, and contained potential errors (e.g., EK100-GT label ‘turn-off tap’ for a video of an actor washing a bowl).

For the purpose of the current study, we defined an *Easy* and *Hard* videos based on MOFO’s classification performance (Ahmadian et al., 2023), which achieves industry leading accuracy on the EK100 of 54.5% (see also Chalk et al., 2024). MOFO’s classification advantage comes from using optical flow to identify a region of significant motion, whose large portions are then masked and subsequently reconstructed in a self-supervised learning process to efficiently learn the spatial and temporal structures used for classification.

We selected 109 *Easy* videos and 128 *Hard* videos. *Easy* videos were defined as videos in which the EK100-GT was the same as the network’s top-1 classification prediction and the confidence of the network was greater than 60%. We defined *Hard* videos as videos in which the EK100-GT was not present among the network’s top-5 most likely classification predictions. We selected a smaller number of *Easy* videos as we found this number sufficient and were able to match the two datasets on key characteristics (e.g., number of consistently recognizable videos per action category).

#### 2.1.1 Video Preprocessing

The selected EK100 videos were processed to produce a base video set of 237 videos. Table 1 presents the statistics of these videos before and after different steps of preprocessing. All videos were temporally and spatially cropped to remove irrelevant information for the recognition of the target action as defined by the EK100-GT label. Temporally, the videos’ start and end frames were selected to correspond with action initiation and termination. Spatially, the videos were cropped to include hands and objects directly involved in the action, and objects judged by the researcher as providing contextual information.

**Table 1.**
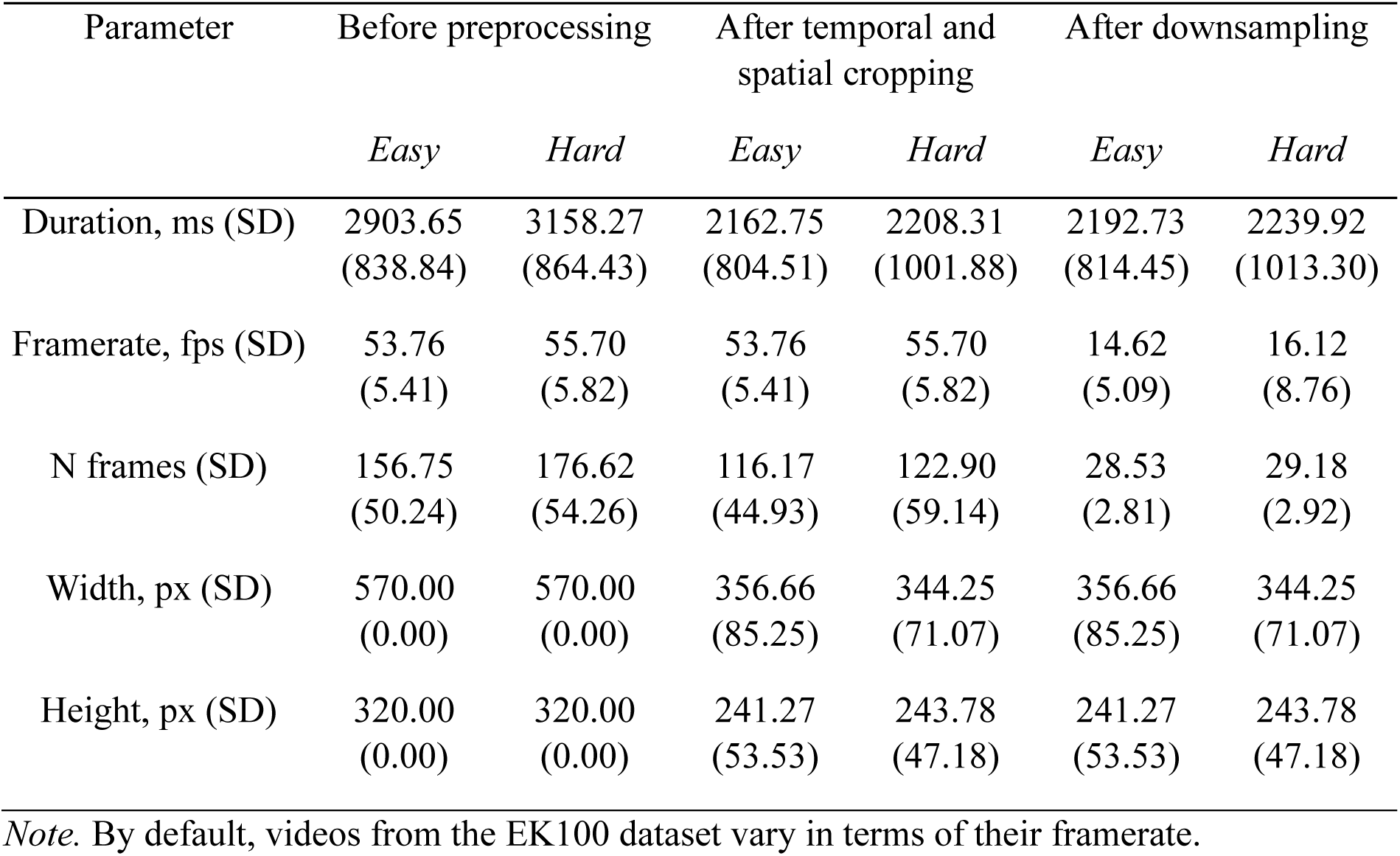
Mean Video Parameters of *Easy* and *Hard* Videos.

| Parameter | Before preprocessing |  | After temporal and spatial cropping |  | After downsampling |  |
| --- | --- | --- | --- | --- | --- | --- |
|  | *Easy* | *Hard* | *Easy* | *Hard* | *Easy* | *Hard* |
| Duration, ms (SD) | 2903.65<br>(838.84) | 3158.27<br>(864.43) | 2162.75<br>(804.51) | 2208.31<br>(1001.88) | 2192.73<br>(814.45) | 2239.92<br>(1013.30) |
| Framerate, fps (SD) | 53.76<br>(5.41) | 55.70<br>(5.82) | 53.76<br>(5.41) | 55.70<br>(5.82) | 14.62<br>(5.09) | 16.12<br>(8.76) |
| N frames (SD) | 156.75<br>(50.24) | 176.62<br>(54.26) | 116.17<br>(44.93) | 122.90<br>(59.14) | 28.53<br>(2.81) | 29.18<br>(2.92) |
| Width, px (SD) | 570.00<br>(0.00) | 570.00<br>(0.00) | 356.66<br>(85.25) | 344.25<br>(71.07) | 356.66<br>(85.25) | 344.25<br>(71.07) |
| Height, px (SD) | 320.00<br>(0.00) | 320.00<br>(0.00) | 241.27<br>(53.53) | 243.78<br>(47.18) | 241.27<br>(53.53) | 243.78<br>(47.18) |

*Note.* By default, videos from the EK100 dataset vary in terms of their framerate.

In addition, for Experiment 1, we downsampled the number of frames by taking every *X*th frame. The number of frames per video was normalized to a fixed target range of 25–35 frames. For each video, the original frame count was linearly scaled from its native range (32–298 frames) to the target range, and the required downsampling factor *X* was calculated accordingly. The frame rate for each video was adjusted to maintain the duration of the base video after spatial and temporal cropping.

### 2.2 Experimental Protocol

The online study for both Experiments 1 and 2 was set up on the Gorilla platform (www.gorilla.sc; Anwyl-Irvine et al., 2020) and disseminated via Prolific (www.prolific.com). The study was approved by the Newcastle University Research Ethics Committee (Ref: 38465/2023). Data collection for Experiment 1 occurred between 27.02.2024 and 19.06.2024.

We tested 136 participants in Experiment 1. Each participant was randomly allocated to label one of two 64-item subsets of *Hard* videos (39 and 37 participants, respectively), a 61-item subset of *Easy* videos, or a 48-item subset of *Easy* videos (30 participants each). Demographic information was lost for nine participants. The remaining participants included 44 females and 83 males, with an average age of 29.9 years old (SD = 20.2 years old). The remuneration for participation was between £2.80 and £4.50, depending on the size of the assigned dataset.

During both online Experiments 1 and 2, participants viewed a series of videos and typed a response to each video. Each trial began with the presentation of a central black fixation cross against a white screen for 500ms. The video was then played in a loop. It was scaled to a height of 200px, on average making up 25.8% of participant’s viewport height (SD = 5.5%), while maintaining the original aspect ratio. After 4000ms, a prompt and a single response box were displayed beneath the video. Participants were instructed to type their response in the form of a single action followed by a single acted-on object, using up to 3 words. After 4 practice trials, participants performed one test trial per video in a randomly presented order. The subset was intermixed with four catch trials, regularly spaced out for the subset. Catch trials used videos from other subsets that were expected to be labelled consistently. No participants were removed from analysis due to poor performance on catch trials. Participants could take an optional 60s break halfway through the session. Full participation in one session took up to 35min.

### 2.3 Behavioral Response Analysis

For each typed response, we calculated its Semantic Similarity to every other response for that video using a custom Python LM framework based on the sentence-BERT (SBERT) model, version ‘all-mpnet-base-v2’ (Reimers & Gurevych, 2019) fetched via the ‘sentence-transformers’ Python package v.2.2.2. Semantic Similarity, *S_sim_*, was computed as follows. After removing punctuation, articles and subjects (words like ‘man’, ‘person’), correcting misspellings algorithmically and using SymSpell (Garbe, 2012), and manual rewording of responses with incorrect word count or slang, pairs of responses were input into SBERT, outputting cosine similarity between their embedding vectors, *CS* (see Figure 1). We also computed the cosine similarity between the isolated actions from the two responses, *CS_A_*, and cosine similarity between isolated objects, *CS_O_*, using Word2Vec (Mikolov et al., 2013), version ‘word2vec-google-news-300’ fetched via the ‘gensim’ Python package v.4.3.0. The *S_sim_* was then calculated as:

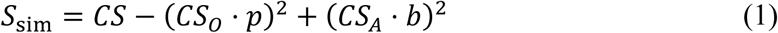

where *p* and *b* are penalty and bonus constants, respectively, optimized a priori. The resulting *S_sim_* ranged from -.102 to 2.891, but equaled 2.750 if the two responses were identical. The approach emphasized the similarity of performed actions and maximized performance of the framework. Thus, description ‘pouring milk’ would be semantically closer to ‘pouring water’ than ‘opening milk’. For some action verb pairs, the framework failed to understand existing differences. Specifically, the issue arose when discriminating verb ‘close’ from ‘open’; ‘move’, ‘put’ and ‘hold’ from each other and from the cluster ‘pick’, ‘take’ and ‘grab’; ‘put on’ from ‘open’ and ‘take off’ from ‘close’; preposition pairs ‘in/out’, ‘up/down’, ‘on/off’. To reflect the dissimilarity of these action responses, action similarity was manually set to 0.1 in 602 *Easy* response pairs (1.29%) and 4780 *Hard* response pairs (3.71%). If the *S_sim_* was greater than threshold *t*, also optimized a priori, the two responses were considered significantly semantically similar to each other.

**Fig. 1.**
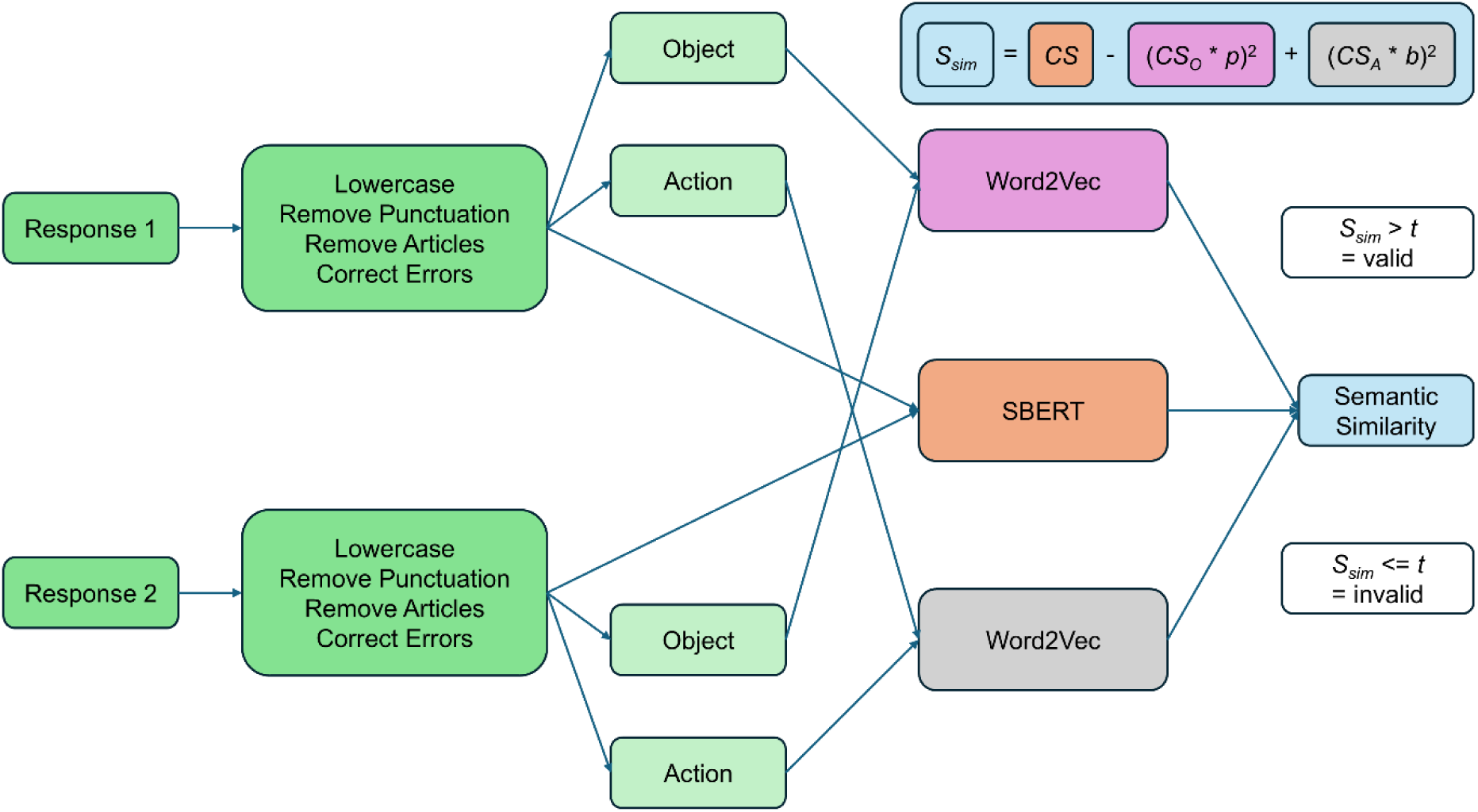
Pipeline for computing semantic similarity between response pairs. Pairs of responses were pre-processed and split into their object and action parts. Full responses and their parts were then input into respective language models to compute cosine similarity between their embeddings. Finally, cosine similarities were input into the formula to calculate Semantic Similarity between responses, *S_sim_*. The formula included penalty (*p*) and bonus (*b*) constants that emphasized similarity of action over object.

#### 2.3.1 Human Ground Truth

For each video, we next determined its HGT response based on the response that was in the centroid of dense neighborhood of semantically similar responses as illustrated in Figure 2. Further visualization of the HGT computation process is presented in Figure S1.1 of the Supplementary material. Once the Semantic Similarity score, *S_sim_*, was calculated for all response pairs for a video, we sorted the scores of each response in descending order. We then took the score at the 75th percentile as the measure of Semantic Neighborhood Density (SND) of that response (Hendrix & Sun, 2021). For instance, if there were 20 responses in total, each response would have 19 Semantic Similarity scores. Sorted in descending order, the 15th score on that list would be the SND for that response. The HGT for a given video was the response with the largest SND.

**Fig. 2.**
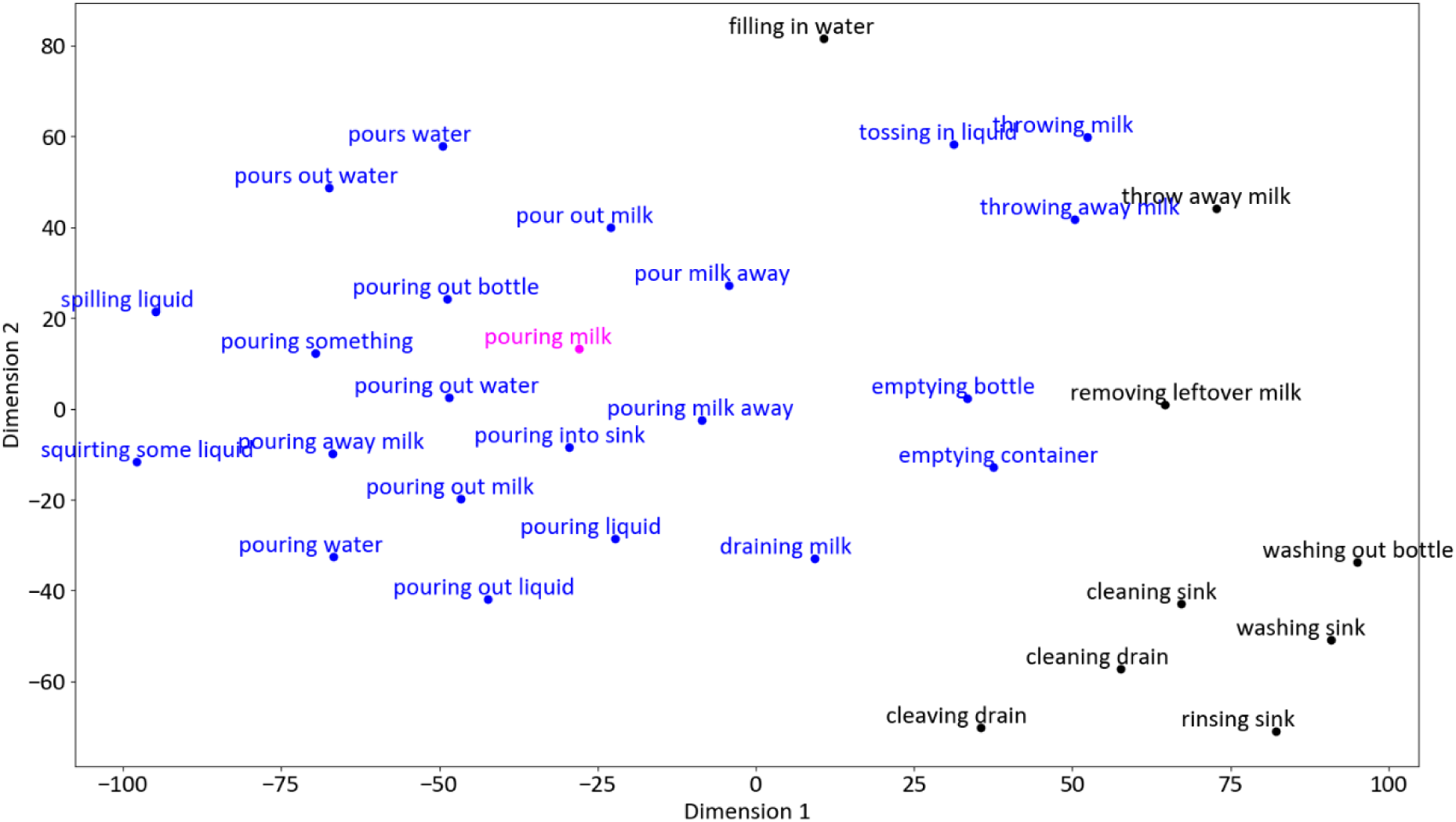
Semantic space for the ‘pouring milk’ video. The HGT (magenta) was defined as the response with the greatest Semantic Neighborhood Density, which means it can fit 75% of semantically closest responses into a minimal space. The semantic distance between the HGT and the other responses presented here is inversely proportional to semantic similarity. Responses in blue were significantly semantically similar to HGT (i.e., *S_sim_* > *t*). Responses in black were not significantly semantically similar to the HGT. This space was constructed through the t-Distributed Stochastic Neighbor Embedding method.

#### 2.3.2 Recognition Consistency

The SND of the selected HGT then served as the measure of Recognition Consistency of the corresponding video. Videos were considered consistently recognizable if their Recognition Consistency was greater than *t* = 0.65 (with *b* = 1.375 and *p* = 0.375). Once a consistent recognizability candidate was highlighted by the SBERT framework, the researcher validated whether its semantically closest responses at the 75th, 65th, 60th, 55th, 50^th^ and 45^th^ percentiles (all with *S_sim_* > 0.65) should be considered correct responses under the selected HGT. Consistent recognizability was confirmed if the researcher agreed with all 6 decisions of the SBERT framework.

### 2.4 Results of Semantic Modelling

Participants viewed short videos of human actions and labelled each video by the action being performed. Mean number of words across all responses after corrections was 2.295 (*SD* = 0.471) for *Easy* videos, 2.335 for *Hard* videos (*SD* = 0.486). Figure 3 presents descriptive statistics from our semantic modelling and compares the semantic similarity of our HGT to the EK100 and MOFO predictions for *Easy* and *Hard* videos.

**Fig. 3.**
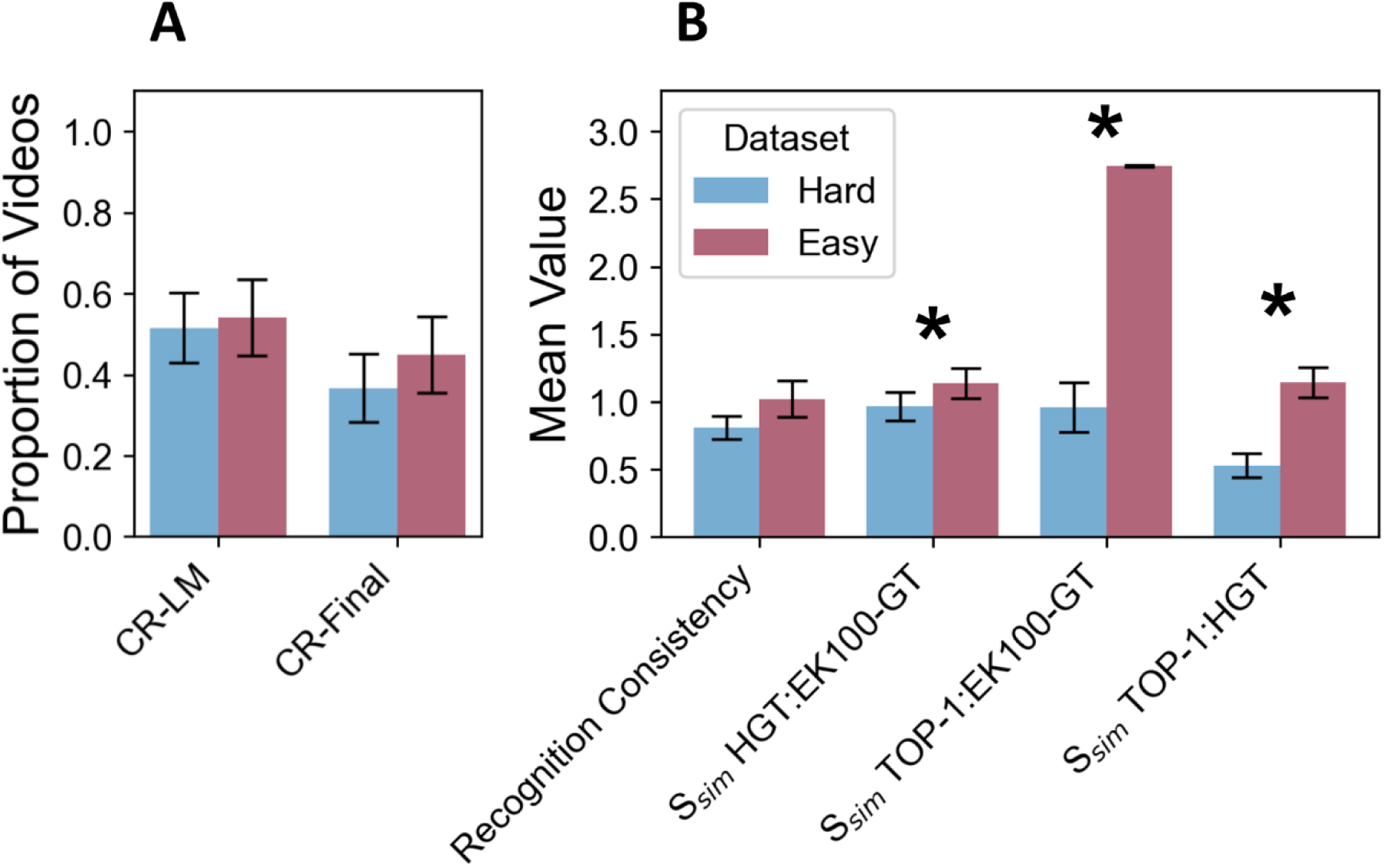
Recognition Consistency results in Experiment 1. A) Mean proportion of *Easy* and *Hard* videos that were consistently recognized (CR-LM = consistent recognizability based on language model framework. CR-Final = consistent recognizability validated by researcher). B) The first bar-pair presents the mean Recognition Consistency of HGT for *Easy* and *Hard* videos. The remaining bar-pairs present the mean Semantic Similarity between the HGT, the EK100-GT and the Top-1 MOFO prediction for *Easy* and *Hard* videos. Error bars reflect 95% confidence intervals. Asterisk ‘*’ indicates a significant difference between *Easy* and *Hard* videos.

#### 2.4.1 HGT Labels and Recognition Consistency

Figure 3A presents the proportion of total videos for which consistent recognizability was signaled by the SBERT framework and the proportion of total videos whose consistent recognizability was also validated by the researcher. The SBERT framework suggested consistent recognizability in 125 of 237 videos, 96 of which were confirmed by the researcher (49 *Easy* videos, 47 *Hard* videos). *Easy* and *Hard* videos did not differ significantly in their probability of being consistently recognizable (non-parametric Chi-Square *χ²* = 1.657, *p* = .198).

The mean Recognition Consistency across all videos was 0.909 (*SD* = 0.611). Recognition Consistency did not differ significantly between *Hard* (*M* = .812, *SD* = .485) and *Easy* (*M* = 1.023, *SD* = .717) videos (non-parametric Mann-Whitney *U* = 6251, *p* = .168).

#### 2.4.2 Semantic Similarity of HGTs, EK100 GTs and MOFO predictions

Figure 3B presents the mean *S_sim_* scores between HGTs and the EK100 GTs and MOFO predictions. HGTs were a valid substitute for EK-100 GTs, developed for compatibility with our LM framework. HGTs for 66.97% of *Easy* videos and 57.03% of *Hard* videos were significantly semantically similar to their corresponding EK100 GT. As expected, top-1 predictions of MOFO were more semantically similar to the EK100 GTs than HGTs (non-parametric Wilcoxon signed-rank *W* = 2250, *p* < .001, *r* = -.838). Top-1 MOFO classification prediction of 8 (6.25%) videos from the *Hard* dataset (3.38% of all) was however significantly semantically similar to the HGT, when it was inaccurate and not significantly similar to the EK100 GT. Nevertheless, consistent recognizability was confirmed for only 3 of the 8, hence HGT was not a superior GT solution than the EK-100 GT for the other 5.

The similarity scores of video labels derived by these different methods differed significantly between the *Easy* and *Hard* videos. *S_sim_* of HGTs to EK100 GTs was greater in *Easy* than *Hard* videos (non-parametric Mann-Whitney *U* = 5810, *p* = .027, *r* = -.167), while *S_sim_* of HGTs to top-1 MOFO predictions was also greater in *Easy* than *Hard* videos (*U* = 2904, *p* < .001, *r* = -.584).

#### 2.4.3 Recognition Consistency of Actions and Objects

To examine the difference in Recognition Consistency of isolated Actions and Objects between *Easy* and *Hard* videos, we isolated each using the SBERT framework, calculated pair-wise *CS_A_* and *CS_o_* between responses for each video and took the *CS_A_* and *CS_o_* at the upper quartile of each action or object as its SND, in the same way as was previously done for full responses. SND of the best performing action and object then served as action and object Recognition Consistency measure for that video.

Figure 4 shows the mean Recognition Consistency as a function of Dataset and Word Role. We conducted a non-parametric 2×2 between-subjects Aligned Rank Transform ANOVA with Recognition Consistency as the dependent variable, and fixed factors Dataset (with 2 levels, *Easy* vs. *Hard*) and Word Role (with 2 levels, Action vs. Object). There was a significant main effect of Dataset (*F*(1, 470) = 6.355, *p* = .012, η² = 0.783), with higher median Recognition consistency for *Easy* videos (*Md* = .332, *IQR* = .277) compared to *Hard* videos (*Md* = .322, *IQR* = .161). The main effect of Word Role (*F*(1, 470) = 6.355, *p* = .254) and the Dataset × Word Role interaction (*F*(1, 470) = 6.355, *p* = .185) were not significant.

**Fig. 4.**
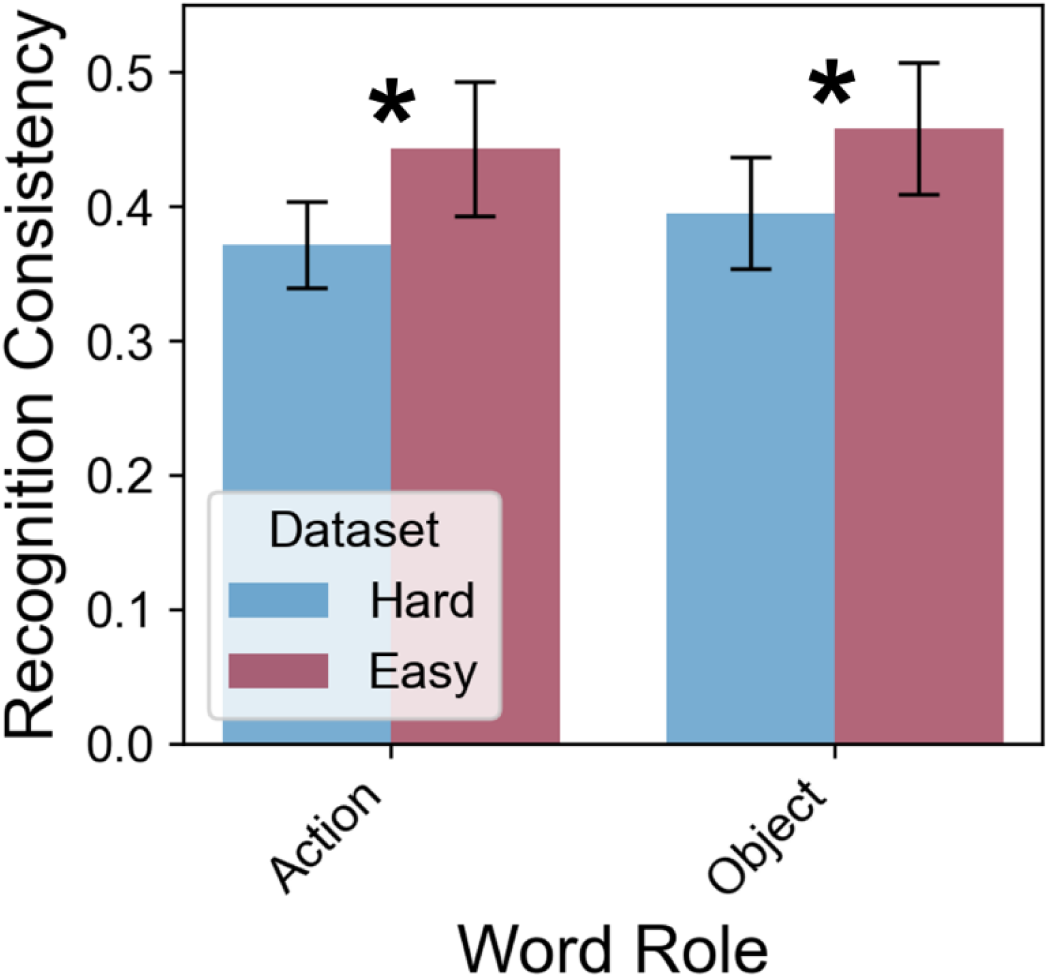
Recognition Consistency for *Easy* and *Hard* videos. Mean Recognition Consistency is presented as a function of Dataset and Word Role. Asterisk ‘*’ indicates a significant difference between *Easy* and *Hard* videos. Error bars reflect 95% confidence intervals.

#### 2.4.4 Clusters of action categories among *Easy* and *Hard* videos

Humans often recognize objects at different levels of abstraction. For example, people can recognize an object as a ‘dog’, rather than a more specific species (‘poodle’) or more general category (‘animal’) (Biederman, 1987). The level of abstraction often depends on observers’ experience with the said object category. Similar levels of abstraction have been shown for allocentric human actions (e.g., ‘to swim backstroke’ vs. ‘locomotion’) (Morris & Murphy, 1990; Zhuang et al., 2023). Therefore, we determined the category structure for the set of 227 egocentric videos tested in Experiment 1, which facilitated stimulus selection for Experiment 2.

To investigate clustering of different action categories among the *Easy* and *Hard* datasets, we calculated pair-wise *S_sim_ between* all *Easy* and *Hard* HGTs and then performed average linkage hierarchical clustering using Python’s scipy library, v.1.11.1 (Rokach & Maimon, 2005), with *S_sim_* when Semantic Distance was equal to zero set to 2.75 (*s*) and clustering distance threshold set to 2.10 (*s* – *t).* Twenty-seven clusters were identified, 20 of which contained HGTs from both datasets. Table 2 presents action category clusters with at least 5 videos in both datasets, while Table S1.1 of the Supplementary Material presents the same data for all 27 identified category clusters. *Easy* and *Hard* datasets did not have major differences in their semantic macrostructure.

**Table 2.**
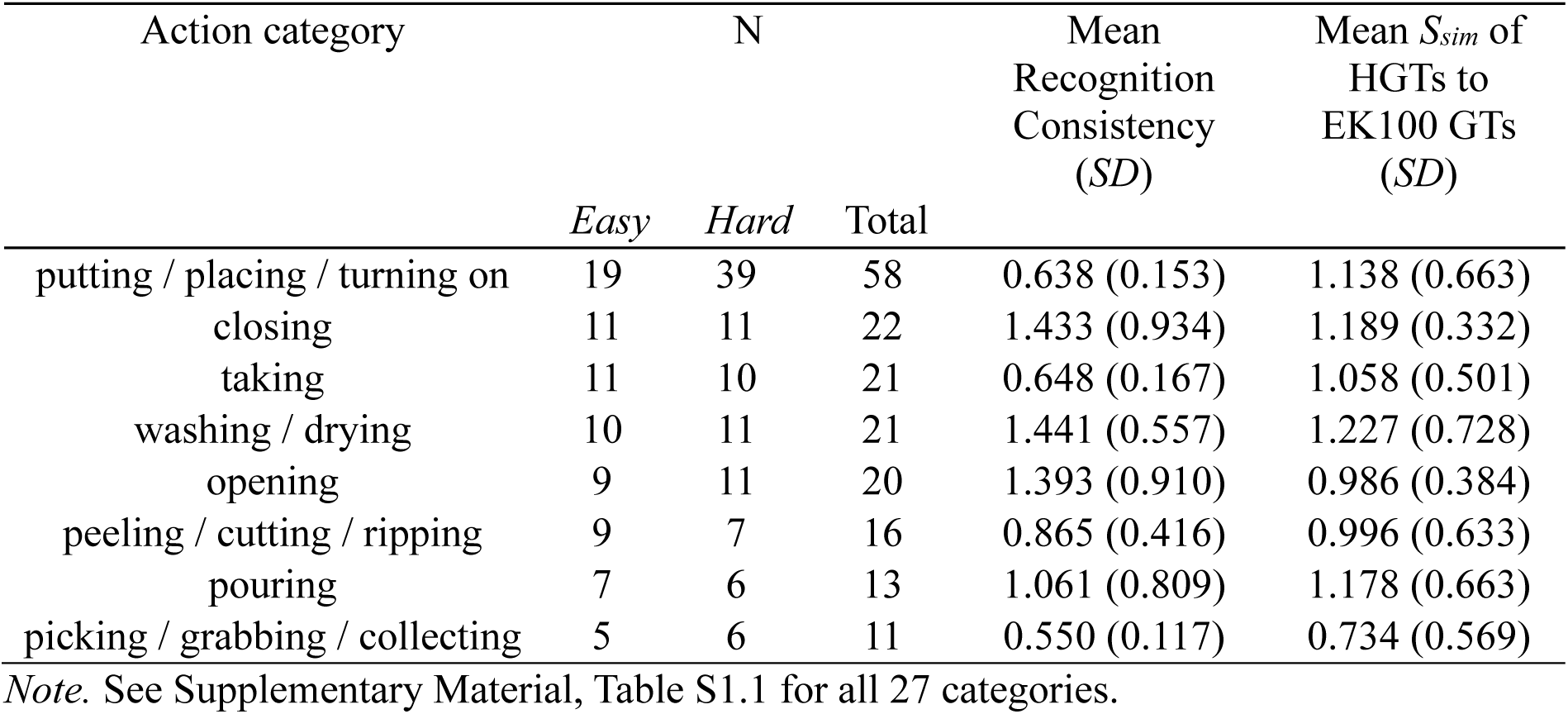
Semantic Clusters of HGTs with at Least 5 Videos in *Easy* and *Hard* Datasets.

## 3. Experiment 2

In Experiment 1, we identified a robust HGT response for 109 *Easy* and 128 *Hard* egocentric action videos from EK100 (Damen et al., 2020). We found that 49 *Easy* and 47 *Hard* videos were consistently recognized (i.e., semantic similarity to the HGT was greater than threshold of 0.65 for 75% of their responses). There were two inter-related aims for Experiment 2. First, we used Ben-Yosef et al.’s (2020) recursive reduction procedure to identify MIRCs for consistently recognized *Easy* and *Hard* videos. Second, we used tree-based classifiers to determine critical mid- to high-level features that are informative for recognizing the actions represented by the MIRCs. Experiment 2 consisted of two phases. In Phase 1, we recursively cropped video frames at each reduction level to identify spatial MIRCs. In Phase 2, we temporally scrambled the frame order of the spatial MIRCs to identify spatiotemporal MIRCs.

### 3.1 Selecting Consistently Recognizable Videos

From the 96 videos identified in Experiment 1 as consistently recognizable, we selected 18 *Hard* and 18 *Easy* videos. We started with three of the HGT categories identified in the first experiment (Table 2): ‘taking’, ‘opening’ and ‘putting / placing / turning on’. For each video type, we selected three videos from each category (9 *Hard*, 9 *Easy*). We then selected *Hard* videos for which the MOFO network’s top-1 classification prediction was ‘take’, ‘open’ or ‘put’ but whose HGT labels belonged to a different action category. There were thus 9 *Hard* false positive videos. Including *Hard* false positives may help us better diagnose and resolve the mistakes that MOFO makes. The remaining *Easy* videos were selected to match the HGT categories of these false positive videos, resulting in 9 *Easy* matching videos. In this experiment, we used the base video set (i.e., no additional frame down-sampling), rescaled to original height of 320px. Tables 3 and 4 present the descriptive statistics for the 36 videos used in Experiment 2 per action category for *Easy* and *Hard* datasets, respectively.

**Table 3.**
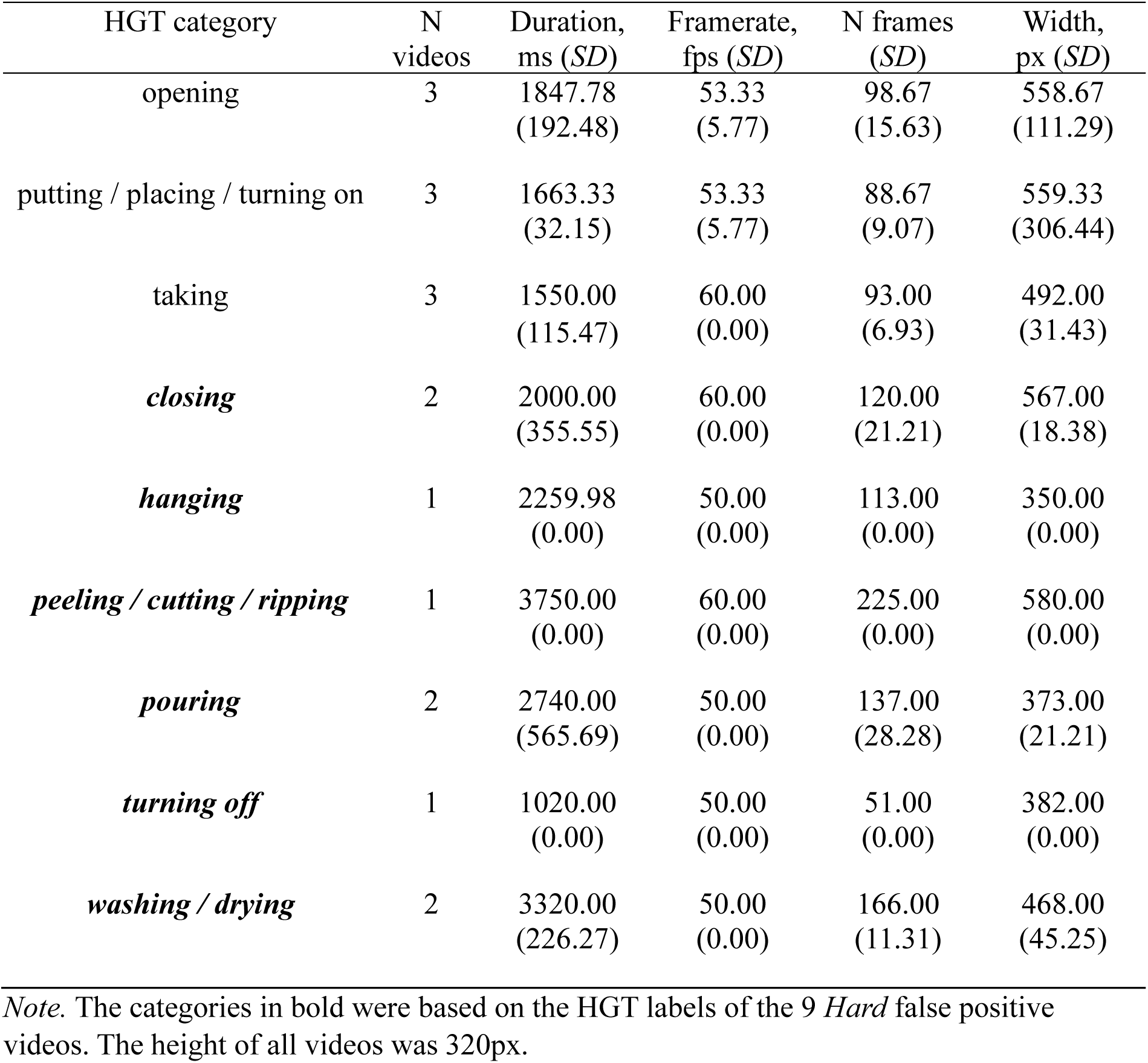
Mean Video Parameters of *Easy* Videos After Preprocessing per Action Category.

| HGT category | N videos | Duration, ms (*SD*) | Framerate, fps (*SD*) | N frames (*SD*) | Width, px (*SD*) |
| --- | --- | --- | --- | --- | --- |
| opening | 3 | 1847.78<br>(192.48) | 53.33<br>(5.77) | 98.67<br>(15.63) | 558.67<br>(111.29) |
| putting / placing / turning on | 3 | 1663.33<br>(32.15) | 53.33<br>(5.77) | 88.67<br>(9.07) | 559.33<br>(306.44) |
| taking | 3 | 1550.00<br>(115.47) | 60.00<br>(0.00) | 93.00<br>(6.93) | 492.00<br>(31.43) |
| ***closing*** | 2 | 2000.00<br>(355.55) | 60.00<br>(0.00) | 120.00<br>(21.21) | 567.00<br>(18.38) |
| ***hanging*** | 1 | 2259.98<br>(0.00) | 50.00<br>(0.00) | 113.00<br>(0.00) | 350.00<br>(0.00) |
| ***peeling / cutting / ripping*** | 1 | 3750.00<br>(0.00) | 60.00<br>(0.00) | 225.00<br>(0.00) | 580.00<br>(0.00) |
| ***pouring*** | 2 | 2740.00<br>(565.69) | 50.00<br>(0.00) | 137.00<br>(28.28) | 373.00<br>(21.21) |
| ***turning off*** | 1 | 1020.00<br>(0.00) | 50.00<br>(0.00) | 51.00<br>(0.00) | 382.00<br>(0.00) |
| ***washing / drying*** | 2 | 3320.00<br>(226.27) | 50.00<br>(0.00) | 166.00<br>(11.31) | 468.00<br>(45.25) |

*Note.* The categories in bold were based on the HGT labels of the 9 *Hard* false positive videos. The height of all videos was 320px.

**Table 4.** Mean Video Parameters of *Hard* Videos After Preprocessing per Action Category.

| HGT category | N videos | Duration, ms (*SD*) | Framerate, fps (*SD*) | N frames (*SD*) | Width, px (*SD*) |
| --- | --- | --- | --- | --- | --- |
| opening | 3 | 2285.56<br>(1621.58) | 56.67<br>(5.77) | 131.00<br>(100.35) | 499.33<br>(54.05) |
| putting / placing / turning on | 3 | 2344.44<br>(1671.44) | 60.00<br>(0.00) | 140.67<br>(100.29) | 317.33<br>(23.09) |
| taking | 3 | 2256.56<br>(1250.59) | 60.00<br>(0.00) | 114.17<br>(53.00) | 473.67<br>(37.64) |
| ***closing***<br>***(‘open’; ‘open’)*** | 2 | 3081.67<br>(1129.01) | 60.00<br>(0.00) | 135.33<br>(75.04) | 455.33<br>(39.31) |
| ***hanging***<br>***(‘insert’)*** | 1 | 933.33<br>(0.00) | 60.00<br>(0.00) | 56.00<br>(0.00) | 538.00<br>(0.00) |
| ***peeling / cutting / ripping***<br>***(‘take’)*** | 1 | 1750.00<br>(0.00) | 60.00<br>(0.00) | 105.00<br>(0.00) | 572.00<br>(0.00) |
| ***pouring***<br>***(‘open’; ‘take’)*** | 2 | 4316.67<br>(141.42) | 60.00<br>(0.00) | 259.00<br>(8.49) | 591.00<br>(55.15) |
| ***turning off***<br>***(‘put’)*** | 1 | 816.67<br>(0.00) | 60.00<br>(0.00) | 49.00<br>(0.00) | 486.00<br>(0.00) |
| ***washing / drying***<br>***(‘take’; ‘put’)*** | 2 | 3586.32<br>(695.35) | 45.00<br>(21.21) | 172.00<br>(108.89) | 440.00<br>(110.31) |

*Note.* The categories in bold were based on the HGT labels of the 9 *Hard* false positive videos. In brackets are inaccurate MOFO top-1 verb predictions for videos in that row. For example, the ‘pouring’ category had 2 videos selected for Experiment 2, one of which MOFO falsely categorized as ‘open’ and one as ‘take’. The height of all videos was 320px.

### 3.2 Experimental Protocol

Data collection occurred between 24.07.2024 and 18.12.2024 for Experiment 2. The first experimental phase of Experiment 2 involved responses from 3800 participants. The second phase involved a further 560 participants (total N = 4360). The sample consisted of 1964 females, 2329 males, 54 non-binary and 13 who preferred not to say, with average age 33.20 years (*SD* = 11.27). The remuneration for participation was between £0.70 and £2.75, depending on the size of the assigned dataset.

#### 3.2.1 Phase 1: Spatial Cropping

For this phase of the experiment, *Easy* and *Hard* videos were cropped into quadrants and tested at different reduction levels in a recursive manner (Ben-Yosef et al., 2020). Figure 5 illustrates this iterative process for a *Hard* video with the HGT response ‘closing container’.

**Fig. 5.**
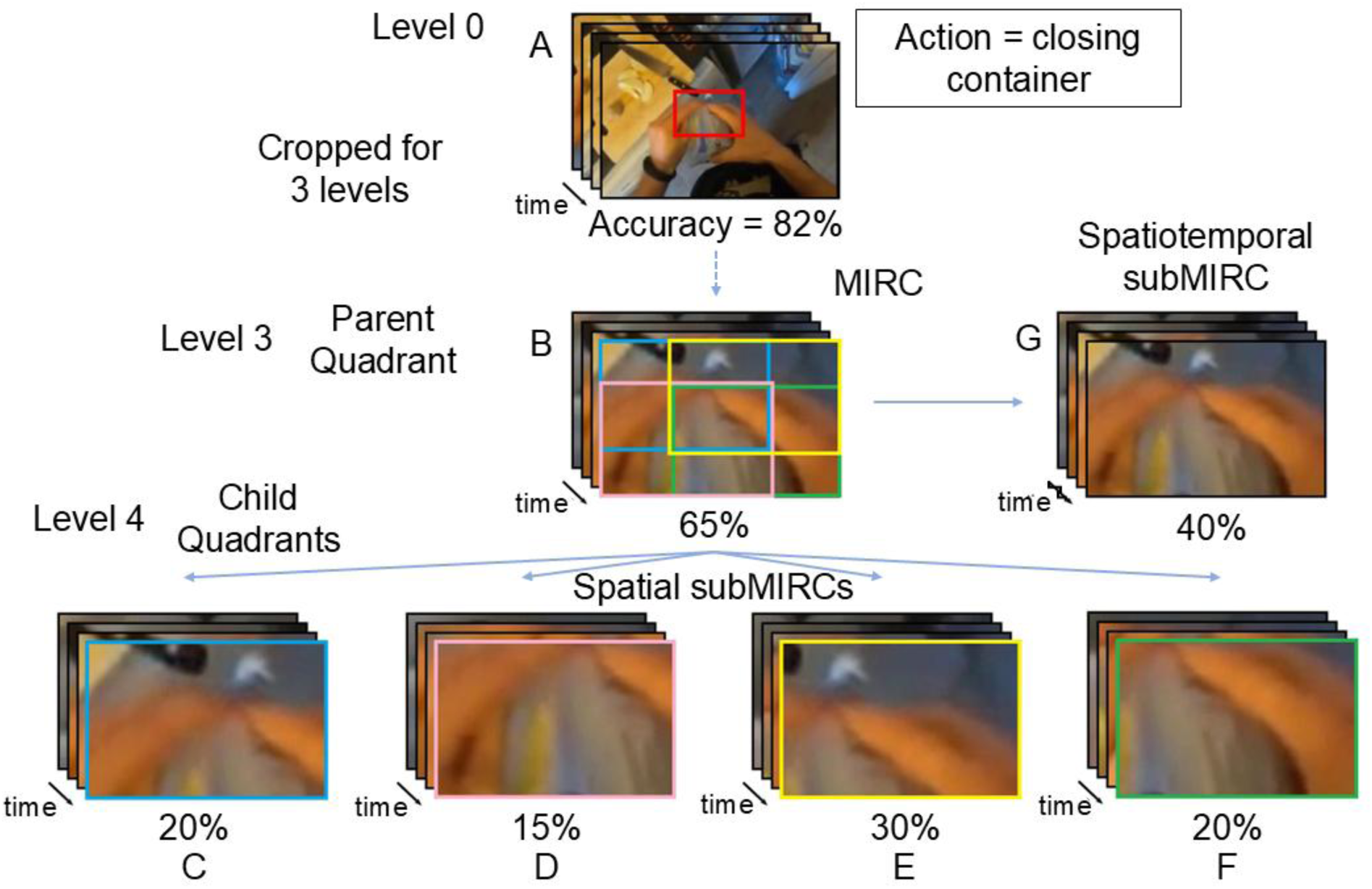
An illustration of the recursive reduction paradigm for the ‘closing container’ video. Recognition responses for the base video A were collected in Experiment 1, where 82% of participants correctly recognized it. In Phase 1 of Experiment 2, A was cropped for 3 Levels before becoming Quadrant B (indicated by red rectangle in A), which was recognized by 65% of participants. Quadrant B served as the Parent Quadrant to produce 4 Child Quadrants at the next reduction level (level 4; C–F). The child quadrants were recognized by *<* 50% of participants, making them subMIRCs and B the spatial MIRC. In Phase 2 of Experiment 2, Quadrant B’s order of frames was scrambled in blocks to create Child Quadrant G, recognized by 40% of participants. Quadrant G is therefore a spatiotemporal subMIRC and Quadrant B is a spatiotemporal MIRC.

We started with the base videos from Experiment 1 (Level 0; Figure 5A). At the first reduction level (Level 1), the 36 videos were cropped into four overlapping quadrants, one at each of their four corners, for a total of 144 quadrants (= 36 × 4). The first reduction removed 40% of the base video frame. Quadrants were considered recognizable if more than 50% of the participants named it correctly (i.e., *S_sim_* to HGT for their response was > threshold 0.65, see Experiment 1; Ben-Yosef et al., 2020). At subsequent reduction levels (i.e., Level 2, 3, 4 and so on), recognizable parent quadrants from the previous level were cropped into smaller child quadrants. These subsequent reductions removed 30% of the parent quadrant. A naive cohort of participants were recruited and tested with the child quadrants, constituting a data collection round. A quadrant was considered a spatial MIRC if all of its four child quadrants at the next immediate reduction level were not recognizable (Figure 5B). These child quadrants were considered as spatial subMIRCs (Figures 5C to 5F). Table 5 presents the breakdown of quadrants as a function of reduction level.

**Table 5.** Number of MIRC and non-MIRC Quadrants per Reduction Level.

| Reduction Level | MIRCs | non-MIRCs |  |
| --- | --- | --- | --- |
|  |  | Recognizable | Unrecognizable |
| 1 | 7 | 123 | 14 (0) |
| 2 | 23 | 317 | 180 (0) |
| 3 | 204 | 627 | 529 (0) |
| 4 | 170 | 721 | 1696 (522) |
| 5 | 69 | 265 | 1890 (1191) |
| 6 | 1 | 0 | 748 (660) |
| 7 | 0 | 0 | 20 (20) |

*Note.* Number of assumed quadrants without a valid recognition accuracy, out of total, is in parentheses.

To avoid a combinatorial explosion of quadrants to test, we created a selection pool of candidate quadrants at each round of data collection. We called this process quadrant pruning. The selection criteria were based on assumptions regarding whether candidate quadrants could be recognized or not. The implications of this selection process were two-fold. First, the recognition status of some quadrants was assumed without testing, based on their overlap with quadrants whose recognition status was known through testing. Thus, we did not measure recognition accuracy for these quadrants. Second, data collection rounds could include quadrants from different reduction levels, as we selected the most informative quadrants from the selection pool and shifted all unselected quadrants into the selection pool for the next data collection round. See Section 3 of the Supplementary Material for more details about the quadrant pruning process.

We recruited enough participants to randomly assign 20 participants to each quadrant at each data collection round. Each participant was tested with only one quadrant from a given video to avoid any repetition effects. Thus, participants were tested with a maximum of 36 quadrants (since there was 36 videos). When there were fewer than 36 videos in a sample set, we ensured that sets matched as best as possible in key properties (e.g., approximately equal number of quadrants from *Easy* and *Hard* videos, from different action categories). This process was iterated until we obtained at least one spatial MIRC for each of the 36 videos. There were seven reduction levels in total, resulting in 7604 total quadrants (474 MIRCs, 1896 subMIRCs, 2053 recognizable non-MIRCs and 3181 unrecognizable non-MIRCs). See Table S3.1 of the Supplementary Material for the impacts of quadrant pruning on composition of the sample sets and Table S3.2 for the distribution of quadrants across data collection rounds. The MIRC and non-MIRC quadrants, as well as their recognition probabilities, are available in the Epic-ReduAct dataset on GitHub (Rahmaniboldaji et al., 2025; https://github.com/SadeghRahmaniB/Epic-ReduAct).

The trial structure in Phases 1 and 2 was the same as in Experiment 1. Each trial began with a 500ms central fixation cross followed by a quadrant video that was looped. The video was scaled to a height of 20.8% of participant’s viewport size, averaging at 170.64px (*SD* = 49.63), while maintaining its aspect ratio. After 4000ms, the response box was presented below the video. Participants could watch the video for as long as they desired and typed what they considered the performed action and the manipulated object, using up to three words. The quadrants were presented in a random order.

The experiment began with five practice trials using simple quadrants of untested, consistently recognized videos from Experiment 1. Participants were then tested with a maximum of 36 test trials. As in Experiment 1, we included two catch trials consisting of easy-to-recognize quadrants with less than three applied reductions, belonging to consistently recognized videos not tested in Experiment 2. These trials occurred at the one-quarter and three-quarter marks of the experimental session. Data from 17 additional participants were not analyzed as they performed poorly on catch trials built into the procedure. There was an optional break half-way through the experiment (up to 60s). There were between 5 and 36 test trials with session durations between 9 and 21 minutes.

#### 3.2.2 Phase 2: Temporal Scrambling

In the second phase, we tested the 474 MIRCs from Phase 1. We temporally scrambled the videos to identify spatiotemporal MIRCs, i.e., MIRCs for which both spatial and temporal information play a role in egocentric action recognition. To avoid large and frequent discontinuities, we scrambled blocks of consecutive frames (e.g., Vuong & Tarr, 2004). Each MIRC video was segmented into five blocks with roughly the same number of consecutive frames. Then the order of these blocks for a given video was randomized with the following constraints: 1) the first block changes position; 2) the last becomes the middle block; and (3) no two originally adjacent blocks remain adjacent. If block scrambling resulted in poor recognition of the MIRC (<50%), then the scrambled quadrant was considered a spatiotemporal subMIRC (Figure 5G) and its MIRC spatiotemporal MIRC. Otherwise, the MIRC remained a spatial MIRC.

We recruited and tested a new cohort of participants. The number of MIRCs varied per video (see Tables S4.1 and S4.2 of the Supplementary Material for the distribution of MIRCs across Easy and Hard videos, respectively). We therefore created different sets of MIRC videos and ensured that sets matched as best as possible on key characteristics (e.g., number of *Easy* and *Hard* videos, videos per action category). Recognition was tested in the same way as in Phase 1. There were 345 spatiotemporal MIRCs (72.78%).

### 3.3 Video Reduction Results

We conducted two main sets of analysis. The first set used linear mixed-effect modelling (LME) to investigate how recognition changed with each level for *Easy* and *Hard* videos. The second analysis set used binary classification to investigate which features can distinguish between different quadrant types (e.g., MIRCs vs unrecognizable non-MIRCs). This will help us test our hypotheses.

#### 3.3.1 Characteristics of MIRCs

Using our reduction methodology, we identified 273 *Easy* and 201 *Hard* MIRCs, with no significant difference in their pixel size (*p* = .383), reduction level depth (*p* = .391) or recognition accuracy (*p* = .603). An average MIRC covered 10210.51 pixels (*SD* = 7608.67), or 3.57% (*SD* = 2.88%) of the area of the original EK100 video, while its center was displaced 112.04 pixels from the center of the original EK100 video, or 17.14% of its diagonal. 200 out of 273 *Easy* MIRCs (73.26%), and 145 out of 201 *Hard* MIRCs (72.14 %) were also spatiotemporal, because their recognition broke down after temporal scrambling.

#### 3.3.2 Metrics of Recognition Decline

We assessed two metrics of recognition decline in *Easy* and *Hard* videos. Reduction rate refers to the difference in recognition accuracy between a child quadrant and its parent quadrant (Rahmaniboldaji et al., 2025) and measures the rate of recognition decline with one level of reduction (which removed 30-40% of the visual information). The reduction rate included cases when the reduction led to a breakdown of reliable recognition (<50%) and when the reduction did not (≥50%). The recognition gap is a special case of the reduction rate and refers to the difference in recognition accuracy between a MIRC (parent) quadrant and its subMIRC (child) quadrants (see Figure 5; Ben-Yosef et al., 2020). The recognition gap also represents a breakdown of reliable recognition. Recognition gap is spatial if between MIRC and its spatial subMIRC, and spatiotemporal if between MIRC and its spatiotemporal subMIRC.

To investigate how the reduction rate changed between *Easy* and *Hard* videos and across reduction levels, we fitted a LME predicting reduction rate from Video Subset (*Easy* vs. *Hard*), the linear effect of Reduction Level (centered), and their interaction, controlling for log Quadrant Area (centered), with random intercepts for video ID and an uncorrelated random slope of Reduction Level by video ID. Models with correlated slopes or factor-coded Reduction Level were singular. For further analyses, we therefore used the uncorrelated random-slope and intercept model, random structure (1 + Reduction Level (centered) || Video ID), which was not singular and had a lower *AIC* than the random-intercept-only model (*ΔAIC* ≈ 31.4).

Figure 6A presents the mean reduction rate as a function of Dataset and Reduction Level. Figure 6B presents the probability distribution of reduction rates at all levels with a 0.1 bin size. The main effect of Dataset (*F*(1, 34.2) = 5.814, *p* = .021, *η²ₚ*= .145) and Reduction Level (*F*(1, 36.0) = 50.658, *p* < .001, *η²ₚ*= .585) were significant. The mean reduction rate was greater in *Hard* (*M* = .284, *SD* = .252) than *Easy* (*M* = .243, *SD* = .239) videos. Moreover, the mean reduction was generally greater at later reduction levels. The Dataset × Reduction Level interaction was not significant (*F*(1, 35.1) = 1.947, *p* = .172, η²ₚ= .053). Thus, rate of recognition decline increased with each further reduction level and was faster in *Hard* than

**Fig. 6.**
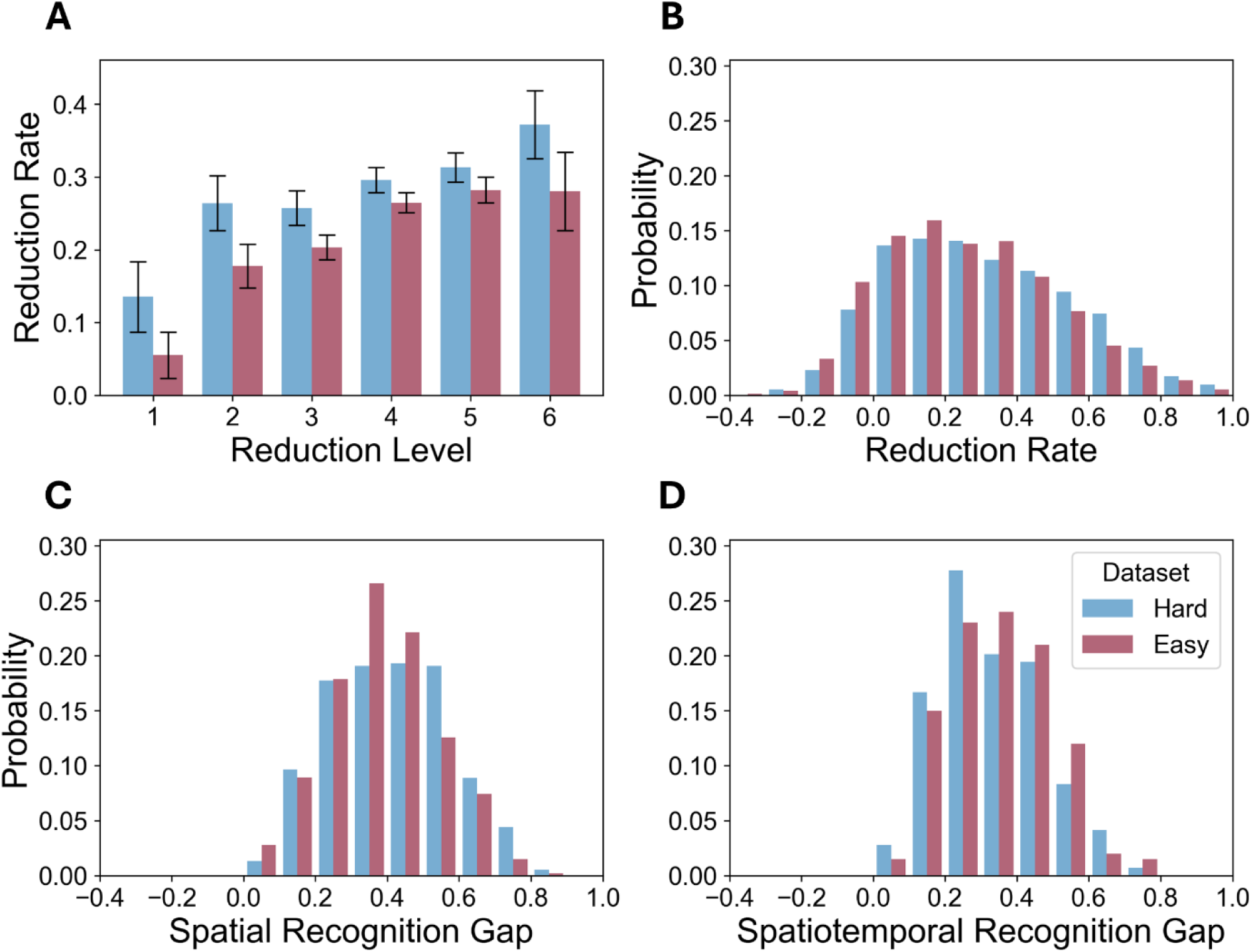
The mean reduction rate and distribution of recognition gap for *Easy* and *Hard* videos in Experiment 2. A) The mean reduction rate at each reduction level for *Easy* and *Hard* videos. Reduction level 7 only involved quadrants with assumed unrecognizability so the reduction rate could not be calculated (see Table 5). Error bars reflect 95% confidence intervals. B) The probability distribution of the reduction rate for *Easy* and *Hard* videos. A negative value meant that recognition improved after the quadrant was reduced. C) The probability distribution of the recognition gap for MIRCs and their Spatial subMIRCs. D) The probability distribution of the recognition gap for spatiotemporal MIRCs and their spatiotemporal subMIRC.

*Easy* videos.

We further examined whether spatial and spatiotemporal recognition gaps differed between *Easy* and *Hard* videos using a Non-parametric Mann-Whitney U test. Figure 6C shows the probability distribution of spatial recognition gap with a 0.1 bin size. There was no significant difference in spatial recognition gap size between *Easy* and *Hard* videos (non-parametric Mann-Whitney *U* = 66568, *p* = .056). The mean spatial recognition gap for *Easy* videos was .358 (*SD* = .151), for *Hard* videos .386 (*SD* = .167). Thus, while on average there was no statistically significant difference in the size of the spatial recognition gaps between *Easy* and *Hard* videos, the distribution peaked marginally lower for *Easy* than *Hard* videos (*p* = .056; see Figure 6C).

Figure 6D shows the probability distribution of spatiotemporal recognition gap with a 0.1 bin size. There was no significant difference in spatiotemporal recognition gap size between *Easy* and *Hard* videos (non-parametric Mann-Whitney *U* = 13538, *p* = .291). Recognition breakdown at the temporal reduction level was thus similar in magnitude between *Easy* and *Hard* videos.

#### 3.3.3 Binary Classification Based on Spatial and Spatiotemporal Features

##### Feature Extraction

We trained random forest binary classifiers to distinguish between different quadrant types based on the following features.

The first set of features quantified the proportions of high-level objects remaining in the quadrant relative to the base video scene, analogous to the Occlusion-Level metric used in amodal segmentation (Qi et al., 2019). We segmented the arms, manipulated objects, and contextual objects, in all frames of the videos using the Segment Anything Model 2 (v.sam2.1_hiera_large; Ravi et al., 2024), run on GPU servers of University of Surrey with 24 GB VRAM. The model output pixel-wise maps for each frame with pixel values belonging to a given object set to 1, otherwise 0. We additionally refined suboptimal masks by hand using a custom Python script. As illustrated in Figure 7, we categorized the segmented objects into the Active Hand (either one or both hands playing the main role in action performance, in contact with the Active Object; Figure 7A), Active Object (up to 2 objects playing the main role in action performance, manipulated by the Active Hand; Figure 7B), Contextual Objects (aggregate area of all other segmented objects potentially having contextual significance, including the forearm of the Active Hand and the passive hand and forearm; Figure 7C) and Background (area not covered by any segmented object).

**Fig. 7.**
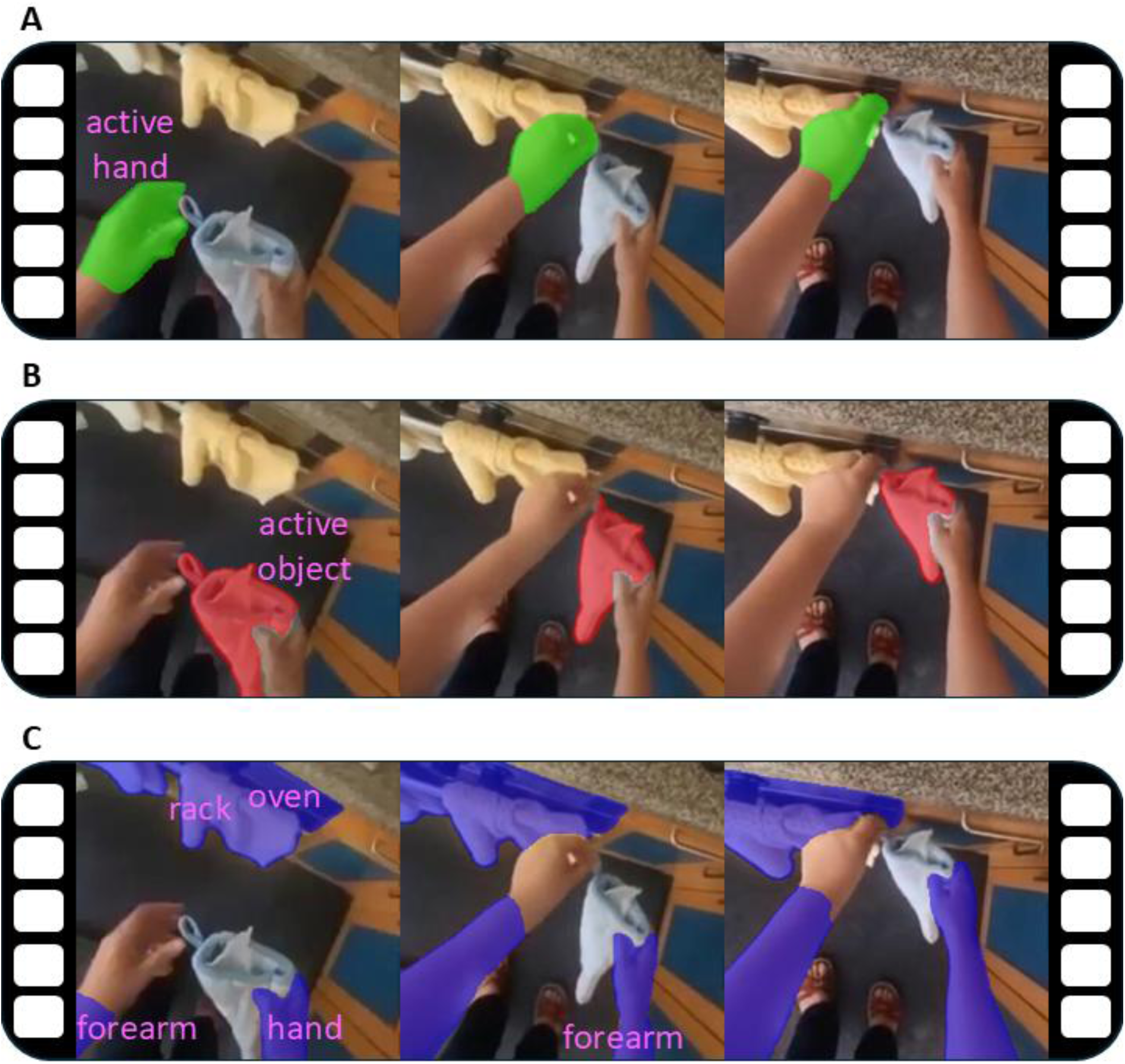
Example segmentations of high-level features related to the surface area of objects in the scene. A) Active Hand, B) Active Object and C) Contextual Objects in the ‘hanging gloves’ video.

The second set of features were the activations of mid-level features in the quadrant relative to the activation of these features in the base video scene. We used the graph-based visual saliency algorithm (GBVS; Harel et al., 2006) to extract conspicuity maps for seven features implemented in the GBVS from the base videos: intensity, orientation, color, flicker, contrast, motion, and two perceptual color spaces. These included the Derrington-Krauskopf-Lennie (DKL) space and the double-opponent (Red-Green/Yellow-Blue) space. Orientation and motion maps were computed for four different directions (0°, 45°, 90° and 135°). These seven conspicuity maps reflect the normalized response between a pixel and its surrounding region for a given feature. Thus, the value at each pixel in the map ranged between 0 and 1, with larger values representing larger activations (relative to the surround) at that pixel. See Section S2 of Supplementary Material for detailed definitions of the mid-level feature activation channels.

We then calculated the proportion of each object’s surface area and each GBVS feature activation score remaining in the frame after cropping. Figure 8 illustrates this process. First, we summed the pixel-wise maps for pixels in the quadrant (*M_q,p_*; red square in Figure 8) and pixels in the full video (*M_f,p_*), across frames (*S_q_*/*S_f_*):

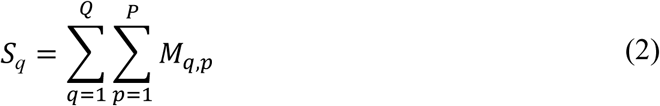

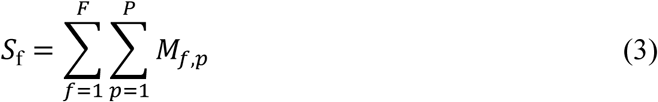

*M_q,p_* and *M_f,p_* represent either the surface area or feature activation. The feature value for each quadrant was then calculated as:

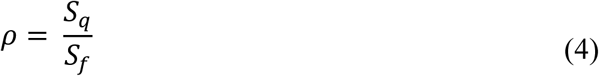

**Fig. 8.**
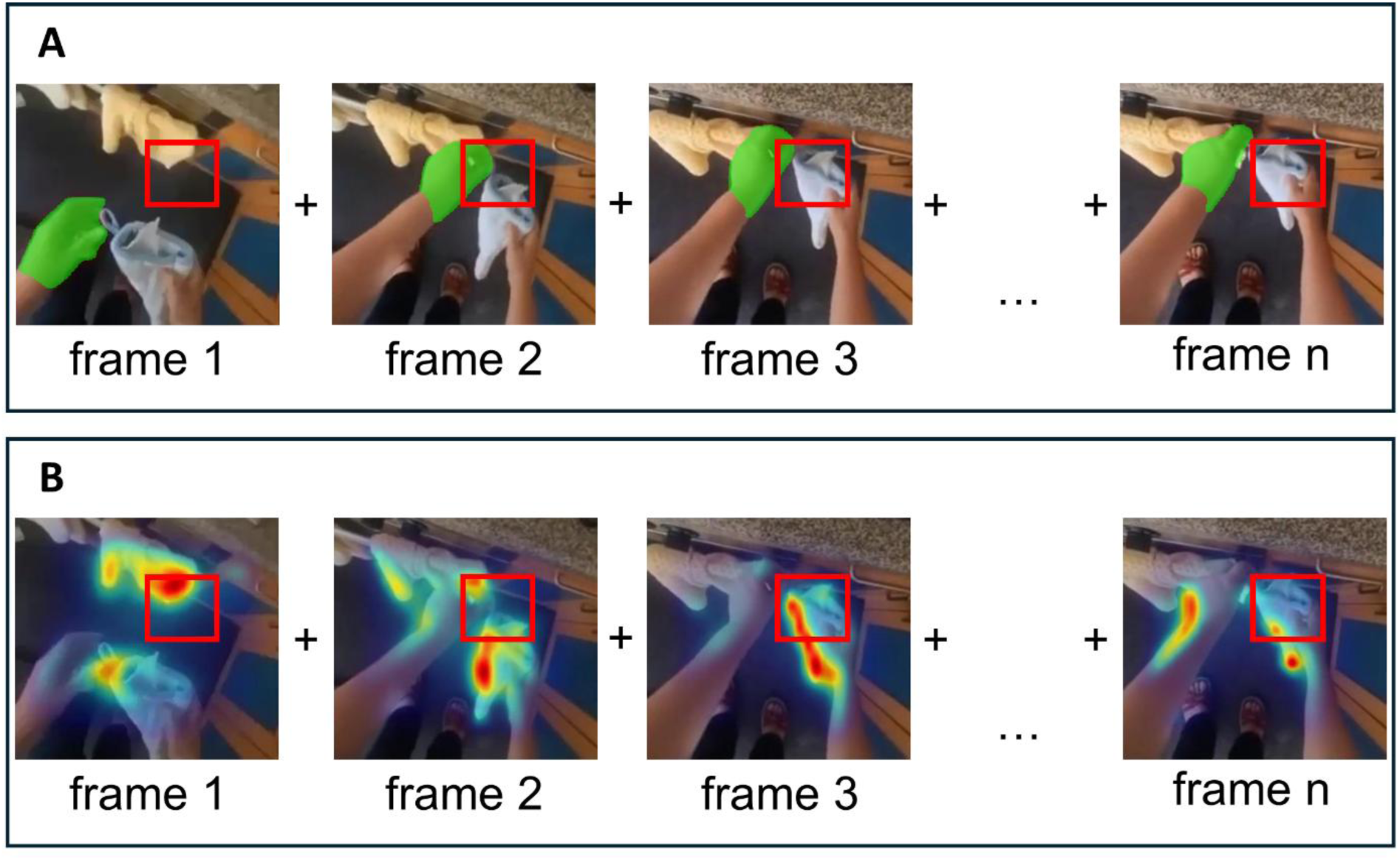
High-level and mid-level feature values within a quadrant for the video ‘hanging gloves’. A) The high-level feature value was based on the mean proportion of pixels belonging to the segmented object contained within the quadrant (red box) relative to the full frame, averaged across all frames. This is illustrated for the Active Hand (green). B) The mid-level feature value was based on the mean proportion of activation of that feature within the quadrant (red box) relative to the full frame, averaged across all frames. The mean activation was computed using GBVS and varied between 0 and 1 (color scale: cool = low activation, warm = high activation). Heatmap in B represents values of conspicuity based on the orientation feature. Example with Active Object and Contextual Objects can be found in Figure S2.1 of the Supplementary Material, while a more detailed depiction of the feature calculation process can be found in Figure S2.2.

##### Classification and Feature Importance Analysis

There were 11 features in total, 4 high-level features based on the visible surface area of objects and 7 based on mid-level visual features. We built random forest binary classifiers, trained and tested either on *Easy* or on *Hard* videos, to allow comparison of the selected features. Our features were skewed and varied over orders of magnitude; therefore, to improve robustness and interpretability of downstream feature importance analysis, we applied a log transform, which does not change the set of candidate partitions in tree-based classifiers. Classifiers were trained using the scikit-learn Python library (scikit-learn v.1.6.1; Pedregosa et al., 2011) running on Python v.3.11.5 and tested using the leave-one-out cross-validation method. Hyperparameters were set to defaults except for the number of estimator trees, which was set to 500 (default 100). A fixed random seed (42) was used for reproducibility. We tested their performance and used SHapley Additive exPlanations (SHAP) technique (Lundberg & Lee, 2018; shap Python library, v.0.45.1) to extract information about feature importance for the classifier. The more positive a SHAP value was, the stronger was the influence of a feature on biasing the model towards classifying a sample as the respective class 1 (e.g., MIRC). In contrast, the more negative a SHAP value was, the stronger was the influence of a feature on biasing the model towards classifying a sample as the respective class 0 (e.g., nonMIRC). The mean absolute SHAP value was used to assess feature importance, the magnitude of each feature’s contribution to the prediction, independent of whether this influence was positive or negative.

We only considered feature importance valid if the classifier’s performance was significantly above chance. This was tested using a one-sided binomial test (Bicego & Mensi, 2023), as well as a permutation test, where the binary category labels *y* were randomly shuffled for 1000 permutations and the classifier was trained and tested on each permutation (Ojala & Garriga, 2010).

We next considered whether individual features predicted quadrant status. This was achieved using the Boruta SHAP method (Kursa & Rudnicki, 2011), which determined whether the feature’s contribution was significantly distinguishable from noise. Boruta was implemented using a custom Python script. We cloned each existing feature (e.g., Active Hand) to produce its shadow feature (Shadow Active Hand) and randomly shuffled the shadow features’ values between videos. We then trained and tested the classifier on the true and shadow features together, to produce their SHAP values. Subsequently, we bootstrapped the SHAP values, with replacement, for 1000 iterations and assessed whether the mean feature importance of true features was higher than the top 95th percentile of the importance distribution of all shadow features. Boruta threshold was used as threshold of significant feature.

##### MIRCs vs. Unrecognizable Quadrants

To determine which mid- and high-level features were critical for recognition, we first compared a set of spatial MIRCs (*Easy* N = 273, *Hard* N = 201) to a subset of unique unrecognized non-MIRC quadrants (*Easy* N = 273, *Hard* N = 201) established in Phase 1, matched as best as possible in number of quadrants per video and their reduction level. Mean reduction level in the MIRC set was 3.58 (*SD* = 0.86), while in the non-MIRC set it was 3.59 (*SD* = 0.83). We used a subset of unrecognized non-MIRCs to maintain class balance, as global feature importance estimates in SHAP are aggregated over the empirical data distribution and can therefore be dominated by the majority class when datasets are highly imbalanced (Lundberg & Lee, 2018), potentially degrading the quality of model interpretation (Chen et al., 2023).

Random forest classifier trained on object surface areas and GBVS feature activation scores to distinguish between *Easy* MIRCs and *Easy* unrecognizable quadrants, achieved accuracy significantly above chance per the binomial test (*Acc* = .634, *p* = .001), with an *F1* score of .638 (*precision* = .631, *recall* = .645). Permutation test also yielded a significant result (*p* < .001), as none of the 1000 random permutations of *y* produced an *F1* score as high as unedited y. In the case of *Hard* MIRCs and unrecognizable quadrants, the classifier also achieved accuracy significantly above chance per the binomial test (*Acc* = .602, *p* < .001), with an *F1* score of .600 (*precision* = .603, *recall* = .597), and the permutation test (*p* = .002), as 1 of the 1000 random permutations of *y* produced an *F1* score as high as unedited y.

Figures 9A and 9B show the mean feature importance and Figures S6.1A and S6.1B of the Supplementary Material present SHAP summary plots, respectively, for *Easy* and *Hard* videos. The SHAP Boruta method yielded significant importance for Active Object and Orientation for both *Easy* and *Hard* videos. In addition, the Background was also significant for *Easy* videos. The Active Hand was not a significant feature for either dataset, contrary to our hypothesis, but interestingly was ranked 4th for *Easy* videos and 7th for *Hard* videos. There was no clear pattern for the ranking of the remaining features.

**Fig. 9.**
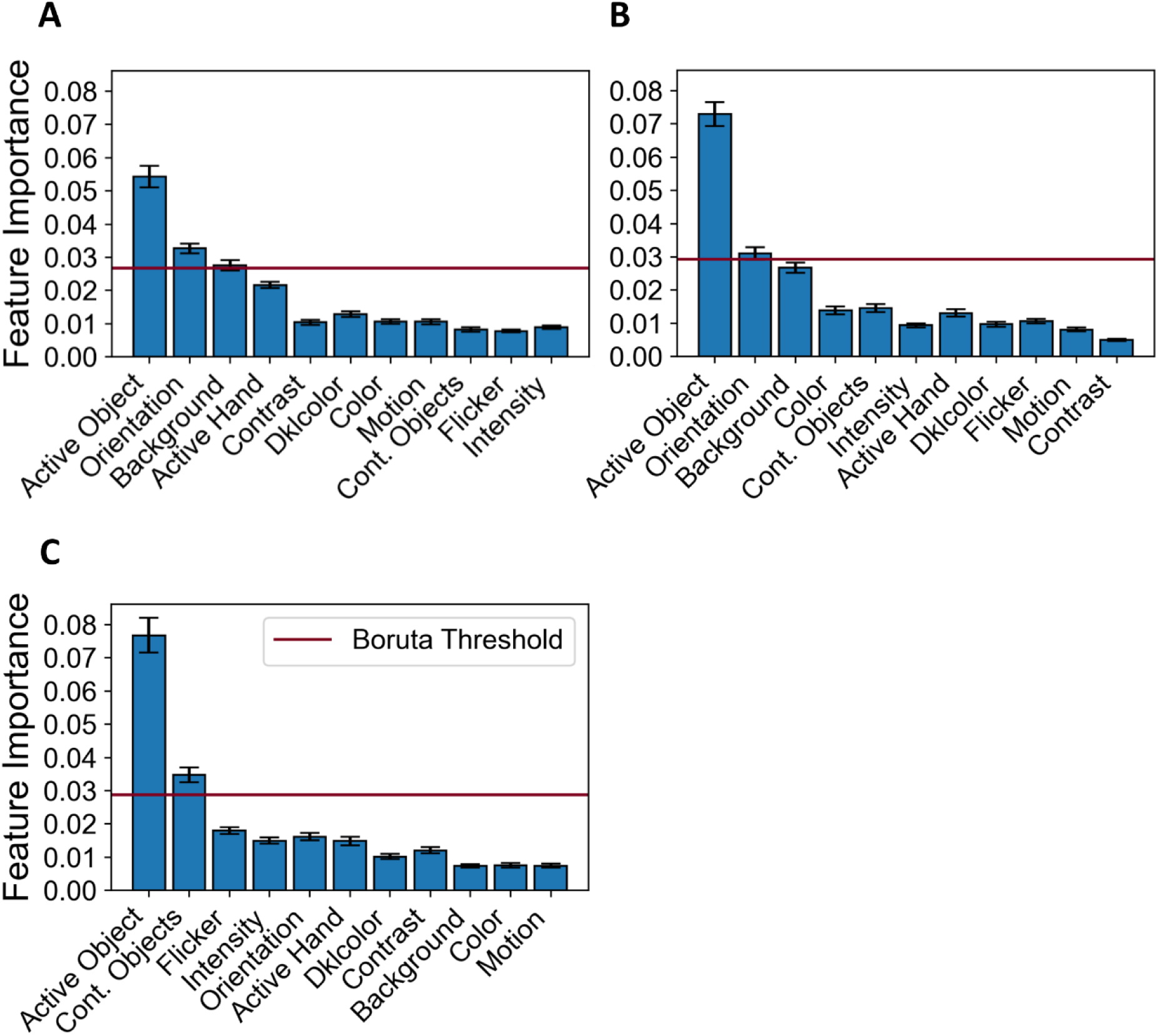
Feature importance for significant binary classifications in Experiment 2. A) Mean feature importance for the classification of MIRC vs unrecognizable quadrants from *Easy* videos. B) Mean feature importance for the classification of MIRC vs unrecognizable quadrants from *Hard* videos. C) Mean feature importance for the classification of *Easy* vs *Hard* MIRCs. Mean feature importance greater than the Boruta Threshold (purple) were significantly informative for the classifier. Features are ordered by their SHAP summaries in Figure S6.1 of the Supplementary Material. Error bars reflect 95% confidence intervals.

Table 6 presents the mean feature values as a function of Quadrant Type and Dataset. When averaged across *Easy* and *Hard* videos, Non-parametric Mann-Whitney U tests showed significant differences between MIRCs and nonMIRCs for all features (*ps* < .025), except Contrast (*p* = .139). In particular, there were larger proportions of Active Object in the MIRC compared to the nonMIRC quadrants (*Easy*: *U* = 27481, *p* < .001, *r* = .263; *Hard*: *U* = 13035, *p* < .001, *r* = .355). There was also higher Orientation activation in the MIRC compared to the nonMIRC quadrants (*Easy*: *U* = 30625, *p* < .001, *r* = .178; *Hard*: *U* = 16241, *p* < .001, *r* = .196). On the other hand, there were larger proportions of Background present in the nonMIRC compared to the MIRC quadrants (*Easy*: *U* = 33415, *p* = .037, *r* = -.103; *Hard*: *U* = 17561, *p* = .024, *r* = -.131). Although Active Hand did not have significant feature importance for our classifier, as initially predicted (Loucks & Baldwin, 2009), this feature was significantly greater in MIRC compared to nonMIRC quadrants (*Easy*: *U* = 32479, *p* = .008, *r* = .128; *Hard*: *U* = 16793, *p* = .003, *r* = .169). Therefore, the visibility of Active Object and Orientation signals was positively associated with recognition, while the visibility of Background was negatively associated with recognition. The other features, including the Active Hand, had negligible impact on classification. See Table 6 for full list of features with indicated significance.

**Table 6.** Mean Feature Values for *Easy* and *Hard* MIRCs and Unrecognizable Quadrants.

| Feature | *Easy* |  | *Hard* |  |
| --- | --- | --- | --- | --- |
|  | MIRC | Unrecognizable | MIRC | Unrecognizable |
| Active Object | **0.124 (0.123)** | **0.078 (0.098)\*** | **0.231 (0.205)** | **0.131 (0.165)\*** |
| Orientation | **0.102 (0.062)** | **0.085 (0.053)\*** | **0.123 (0.083)** | **0.100 (0.076)\*** |
| Background | **0.044 (0.052)** | **0.053 (0.056)\*** | 0.050 (0.072) | 0.056 (0.061)* |
| Active Hand | 0.101 (0.152) | 0.073 (0.122)* | 0.146 (0.183) | 0.102 (0.153)* |
| Color | 0.081 (0.055) | 0.072 (0.049)* | 0.093 (0.081) | 0.082 (0.071)* |
| Contextual Objects | 0.070 (0.067) | 0.066 (0.065) | 0.109 (0.124) | 0.093 (0.122)* |
| DKLcolor | 0.085 (0.058) | 0.074 (0.049)* | 0.099 (0.075) | 0.086 (0.063) |
| Intensity | 0.087 (0.053) | 0.074 (0.042)* | 0.092 (0.074) | 0.085 (0.063) |
| Contrast | 0.084 (0.061) | 0.077 (0.052) | 0.089 (0.086) | 0.082 (0.076) |
| Motion | 0.085 (0.054) | 0.075 (0.048)* | 0.104 (0.082) | 0.089 (0.067) |
| Flicker | 0.082 (0.049) | 0.074 (0.042) | 0.101 (0.075) | 0.089 (0.067) |

*Note.* *SD* is in parenthesis. Bold values indicate significantly informative features for the binary classification MIRC vs unrecognizable quadrants within a given dataset. Asterisk “\*” indicates a significant difference in mean feature value between MIRC and unrecognizable quadrants within the given dataset.

##### Spatiotemporal MIRCs vs. Spatial MIRCs

We found that visibility of Active Object and Orientation signals was critical for recognition. To examine whether the motion in regions occupied by the 11 features was critical for recognition, we next determined whether they could discriminate MIRCs whose recognizability in Phase 2 broke down with temporal scrambling and MIRCs whose recognizability did not. We thus compared a set of spatial MIRCs (*Easy* N = 73, *Hard* N = 56) to a subset of spatiotemporal MIRCs (*Easy* N = 73, *Hard* N = 56), matched as best as possible in number of quadrants per video and their reduction level. Mean reduction level in both sets was 3.60 (recognized, *SD* = .70; unrecognized, *SD* = .85).

We built random forest binary classifiers trained on object surface areas and GBVS feature activation scores of *Easy* or *Hard* MIRCs. Classifier trained on *Easy* MIRCs failed to achieve accuracy significantly above chance per the binomial test (*Acc* = .514, *p* = .402), with an *F1* score of .530 (*precision* = .513, *recall* = .548). Permutation test also yielded a non-significant result (*p* = .300), as 249 of the 1000 random permutations of *y* produced an *F1* score higher than the unedited *y*. Classifier trained and tested on *Hard* MIRCs also failed to achieve accuracy significantly above chance per the binomial test (*Acc* = .563, *p* = .110), with an *F1* score of .559 (*precision* = .564, *recall* = .554) and the permutation test (*p* = .093), as 67 of the 1000 random permutations of *y* produced an *F1* score higher than the unedited *y*. These results indicate that the critical motion in spatiotemporal MIRCs wasn’t reliably tied to any of our defined features. Thus, we did not investigate feature importance further; however, we chart feature importance in Figure S5.1 of the Supplementary Material, while SHAP summaries are presented in Figures S6.1C and S6.1D, for *Easy* and *Hard* videos, respectively. Table S5.1 of the Supplementary Material further presents the mean feature values and significant contrasts as a function of MIRC Type and Dataset.

##### Easy vs. Hard MIRCs

We found that both the high-level Active Object and mid-level Orientation features could predict recognizable MIRC quadrants from unrecognizable quadrants of similar sizes and depth of reduction. There were no reliable features that could discriminate between spatial and spatiotemporal MIRCs and thus represent the sources of critical motion. Furthermore, there was evidence that the contribution of several of our selected mid- and high-level features to egocentric action recognition may differ between *Easy* and *Hard* videos, including the Active Hand (see Figure 9C). There were also some differences affected by temporal scrambling.

Thus, we constructed a classifier to determine features that directly discriminated MIRCs from *Easy* and *Hard* videos. We compared *Hard* MIRCs (N = 201) and a subset of *Easy* MIRCs (N = 201 out of 273) matched as best as possible on their reduction level and HGT action category. Reduction level means of the sample sets were 3.53 (*SD* = 0.93) and 3.56 (*SD* = .85), respectively.

A single random forest binary classifier, trained on object surface areas and GBVS feature activation scores to distinguish between *Easy* and *Hard* MIRCs, achieved accuracy significantly above chance per the binomial test (*Acc* = .761, *p* < .001), with an *F1* score of .766 (precision = .751, recall = .781), and the Permutation test (*p* = .001), as none of the 1000 random permutations of *y* produced an *F1* score higher than the unedited *y*.

Figure 9C shows the mean feature importance plot and Figure S6.1E of the Supplementary Material presents the SHAP summary plot for this analysis. Table 7 presents the mean feature values for *Easy* and *Hard* MIRCs. Only Active Object and Contextual Objects were significantly more informative features than noise. There were significantly larger proportions of Active Object (non-parametric Mann-Whitney *U* = 12295, *p* < .001, *r* = .391) and Contextual Objects (*U* = 17011, *p* = .006, *r* = .158) in the *Hard* compared to the *Easy* MIRCs. There was also significantly more Active Hand (*U* = 17364, *p* = .014, *r* = .140) and Flicker (*U* = 17841, *p* = .043, *r* = .117) in the *Hard* MIRCs, but these features were not significantly informative for our classifier. Thus, *Hard* MIRCs were characterized by including more Active Object and Contextual Objects than *Easy* MIRCs.

**Table 7.** Mean Feature Values for *Easy* and *Hard* MIRCs.

| Feature | *Easy* MIRC | *Hard* MIRC |
| --- | --- | --- |
| Active Object | **0.109 (0.115)** | **0.231 (0.205)\*** |
| Contextual Object | **0.073 (0.073)** | **0.109 (0.124)\*** |
| Flicker | 0.085 (0.053) | 0.101 (0.075)* |
| Intensity | 0.090 (0.059) | 0.092 (0.074) |
| Orientation | 0.106 (0.065) | 0.123 (0.083) |
| Active Hand | 0.117 (0.167) | 0.146 (0.183)* |
| DKLcolor | 0.090 (0.064) | 0.099 (0.075) |
| Contrast | 0.086 (0.064) | 0.089 (0.086) |
| Background | 0.047 (0.055) | 0.050 (0.072) |
| Color | 0.085 (0.058) | 0.093 (0.081) |
| Motion | 0.088 (0.059) | 0.104 (0.082) |

*Note.* *SD* is in parenthesis. Bold values indicate significantly informative features for the binary classification *Easy* vs *Hard* MIRC within a given dataset. Asterisk ‘\*’ indicates a significant difference in mean feature value between *Easy* and *Hard* MIRC.

## 4. Discussion

For human observers, action recognition is an everyday task that is performed quickly, accurately and without conscious effort. By comparison, computer-vision networks only achieve classification accuracy of ∼54.5% on the EK100 dataset (e.g., Ahmadian et al., 2023; Chalk et al., 2024). One key difference between human and computer vision are the visual features that are represented for action recognition. Here we used a new combination of a language model (LM), machine learning and experimental paradigms to identify critical features for recognizing home activities from a challenging egocentric perspective.

In Experiment 1, we identified robust Human Ground Truth (HGT) responses based on semantic similarity for egocentric action videos from the Epic-Kitchens-100 dataset (EK100; Daman et al., 2020). We further identified *Easy* and *Hard* videos which were consistently recognized by human observers. *Easy* videos were ones which the MOFO network (Ahmadian et al., 2023) classified accurately, whereas *Hard* videos were ones which the network failed to recognize. Interestingly, human observers showed greater variability (lower consistency) in some aspects of recognizing *Hard* compared to *Easy* videos.

In Experiment 2, we focused on finding critical mid- to high-level features that can be used to consistently recognize a subset of the *Easy* and *Hard* videos from Experiment 1. For this purpose, we combined our LM with a recursive reduction procedure (Ben-Yosef et al., 2020) to find minimal recognizable configurations (MIRCs). We identified spatial MIRCs, and showed that they are centered around active objects and regions where orientation of edges stands out from the background. We also demonstrated that the MIRCs of *Hard* videos are more likely to involve the active and contextual objects than MIRCs of *Easy* videos. Lastly, we identified spatiotemporal MIRCs by scrambling the frame order of spatial MIRCs. However, none of the critical features tested could discriminate between spatial and spatiotemporal MIRCs. In sum, we identified both mid-level (e.g., orientation) and high-level (e.g., active hand or active objects) features that are critical for egocentric action recognition. These features were predominantly spatial, rather than spatiotemporal features. In the next sections, we discuss the implications of our findings for understanding visual features and hierarchical action recognition, and for action classification in computer vision.

### 4.1 Recognition Consistency of Egocentric Actions

In Experiment 1, we found that 96 of the 237 videos (40.1%) selected from the EK100 dataset were consistently recognized—that is, these videos had a high density of semantically similar responses across participants. Broken down further, this constitutes 49 out of 109 *Easy* (45.0%) and 47 out of 128 *Hard* (36.7%) videos. Low semantic consistency was also found in Smekal et al.’s (2024) study with 10 allocentric mimed action videos. They reported that only 42.1% of responses per video were semantically consistent.

There was no difference in recognition consistency between *Easy* and *Hard* videos when considering the full response. However, *Easy* videos had a significantly higher recognition consistency than *Hard* videos for the action and the object component of the full response. Furthermore, *Easy* video responses had higher semantic similarity to the EK100 GT labels. A related concept in the literature to our recognition consistency is name agreement, which can be expressed as the percentage of participants providing the most frequent name (e.g., for picture naming) or an entropy statistic that takes into account the total number of unique labels for a given item (e.g., Cuetos & Alija, 2003; Perret & Bonin, 2019). These studies suggest that name agreement is associated with visual factors (e.g., body posture, viewpoint or occlusion), motor content and imaginability, and conceptual and linguistic properties. Our findings suggest that *Hard* videos may have a lower name agreement than *Easy* videos for actions. Future studies can thus investigate which of these factors potentially mediate differences in recognition consistency for *Easy* and *Hard* videos.

### 4.2 Spatial and Spatiotemporal Features of Action Recognition

In Experiment 2, we showed that human recognition of actions involving hand-object interactions from an egocentric perspective is predominantly driven by the visibility of the actively manipulated object and the strength of orientation signals in the scene for both *Easy* and *Hard* videos. The absence of scene background also contributed to egocentric action recognition for *Easy* videos. These findings are consistent with our hypothesis that localized information in the vicinity of the hand is important for action recognition in general (see also Loucks & Baldwin, 2009; Ben-Yosef & Ullman, 2018). Furthermore, the recognition of active objects in the scene can help constrain the action classification search space in line with the reverse hierarchy theory (e.g., Hochstein & Ahissar, 2002; Bar, 2003).

We did not find that the active hand was a critical high-level feature for egocentric action recognition, in contrast to our hypothesis and the suggestion of Loucks and Baldwin (2009). One possible explanation lies in our feature extraction procedure. Specifically, we did not differentiate between the fingers and the wrist when quantifying active hand surface area, despite their form and purpose varying systematically. There are however also broader methodological differences between our study and Loucks and Baldwin’s. First, their study presented action videos from an allocentric perspective. Differences between egocentric and allocentric perspective can lead to greater attention on active objects (e.g., Borghi et al., 2012) or on the actor’s body posture including their hands (e.g., Henderson et al., 2008). Second, Loucks and Baldwin did not spatially reduce their videos and therefore did not converge on the minimal features required for recognition. More generally, while the presence of active objects may be specific to actions and efficiently constrain classification (Hochstein & Ahissar, 2002; Bar, 2003), the presence of active hands is consistent across all manipulative actions. Observers must therefore analyze the active hand’s shape and motion, requiring greater cognitive capacity and a larger amount of spatiotemporal information. While it may therefore be less practical to use the active hand for naturalistic action recognition, our findings only imply that the spatiotemporally most limited informative feature configurations (i.e., MIRCs) are less likely to include the active hand than the object. Future studies could triangulate on whether the active object dominance generalizes to full videos, for example with the use of eye-tracking.

As mentioned above, of the mid-level features, only the strength of orientation signals in the scene contributed significantly to egocentric action recognition. This finding contradicts our hypothesis that egocentric action recognition would be dominated by motion signals (Kooiker et al., 2016; Smekal et al., 2024). However, our findings are consistent with the visual processing hierarchy: edges are used to form mid-level contour and curvature features (Pasupathy & Connor, 2001; 2002); the contours, in turn, can help segment high-level objects and object parts (Marr, 1982); and lastly, segmented objects can lead to conceptual representations of the scene (Biederman, 1987). Furthermore, distinctly oriented edges often occur at contour contacts, junctions and corners and may have higher density in ‘small-scale envelopes’, like hand-object manipulations, which depend on fine-grained features such as finger position (Ben-Yosef & Ullman, 2018; Tarhan & Konkle, 2020). The findings are also consistent with visual search (Rutishauser & Koch, 2007) and eye-tracking studies (Suda & Kitazawa, 2015; Kojovic et al., 2024).

We showed that egocentric action recognition may be dominated by spatial features. There was evidence, however, that temporal and/or spatiotemporal features may contribute to recognition. In Experiment 2, we found that scrambling the frame order led to a recognition breakdown of *Easy* and *Hard* spatial MIRCs (see also Vuong & Tarr, 2004; Smekal et al., 2024). However, we did not identify any critical features that significantly discriminated spatial from spatiotemporal MIRCs, which is not consistent with our hypothesis.

### 4.3 Spatial and Spatiotemporal Features in *Easy* vs *Hard* Videos

Lastly, we compared differences between *Easy* and *Hard* MIRCs in Experiment 2, as these differences have further implications for computer-vision systems. We found that recognition decreased at a faster rate for *Hard* than *Easy* videos in Experiment 2 (i.e., greater mean reduction rate at all reduction levels). This finding suggests that removing a random region from *Hard* videos reduced its recognition to a greater degree than removing an equal sized region from *Easy* videos, even though the two video types have equivalent recognition consistency and MIRC recognition probability. In the special case of the recognition gap, which reflects a breakdown of reliable recognition between MIRCs and all their children, we found an overall mean gap of .37 (*SD* = .16) and a marginally significant difference between *Easy* and *Hard* videos (*p* = .056). Our mean recognition gap is comparable to Ben-Yosef et al.’s (2020) gap of ∼.43 with allocentric actions (e.g., playing a violin, rowing, mopping) and Ullman et al.’s (2016) gap of ∼.57 with static images of objects (e.g., airplane, eagle, horse). Observed cross-experiment differences may depend on the complexity and availability of different spatial and spatiotemporal features. For example, increasing complexity may introduce more high-level features which can substitute the informativeness of one another and lead to smaller recognition gaps.

We also found that the active object and contextual objects discriminated *Easy* from *Hard* MIRCs when we directly tested these two MIRC types for all features. The importance of orientation (and background for *Easy* videos) did not significantly differentiate between the two MIRC types, even though this feature discriminated MIRCs from unrecognizable quadrants. Thus, compared to computer vision, human observers may rely on active objects and high-level context (e.g., other objects in the scene) to recognize *Hard* videos with the same consistency and accuracy as *Easy* videos, in line with the reverse hierarchy framework (Ahissar & Hochstein, 2004; Hochstein & Ahissar, 2002).

## 5. Conclusions and Implications for Computer Vision

In the current study, we combined a language model with a recursive reduction paradigm to identify critical features that are informative for action recognition. The language model allowed us to find robust ground truth labels for a variety of actions carried out in a home environment, taking into account the variability of human responses. Importantly, the human ground truth labels were used to identify minimal configurations and within these, informative features for action recognition. These features included the active object involved in the action and orientation signals in the scene. Although we focused on recognizing egocentric actions from a home environment, our approach of identifying mid- (e.g., orientation) to high-level (e.g., hands and objects) features can generalize to other types of actions or other challenging viewing conditions (e.g., poor-resolution videos or cluttered environments).

State-of-the-art wearable devices for AI-driven task assistance recognize objects, locations, or predefined cues, but fail to dynamically understand the structure of ongoing actions (e.g., Huh et al., 2025; Vuzix Corporation, 2026). In line with these limitations, while human observers were able to recognize all actions in our experiments, computer-vision networks failed to recognize some of the actions (i.e., the *Hard* videos). However, we found differences in how the different features contributed to the recognition of *Easy* and *Hard* videos for human observers. Given these differences, the spatial and spatiotemporal MIRCs can be used to construct more human-like computer-vision networks and models for action recognition (Ben-Yosef et al., 2018; Ben-Yosef et al., 2020; Rahmaniboldaji et al., 2025). For example, these models could focus their feature extraction on MIRC regions (Ahmadian et al., 2023; Tong et al., 2022) to improve the classification of *Hard* videos. Such human-inspired models can, thus, perform better at complex tasks and be more efficient, compact and interpretable (Kubilius et al., 2019). In sum, integrating features used by humans into artificial networks can allow incorporation of higher-order human-like processes currently absent in artificial systems (Chrol-Cannon et al., 2021).

## Data Availability

Data is available through GitHub (https://github.com/filip1O/Seeing-Just-Enough-The-Contribution-of-Visual-Features-to-Egocentric-Action-Recognition).

