## Supplemental Materials 1 for "Seeing Just Enough: The Contribution of Hands, Objects and Visual Features to Egocentric Action Recognition"

**S1. Human Ground Truth Calculation and Action Categories from Experiment 1**

**A**

| Comparison Response | Pouring Water | Washing Sink | Pouring Milk | … | Emptying Bottle |
|---|---|---|---|---|---|
| Pouring Water | 2.75 | 0.42 | 2.61 | … | 0.80 |
| Washing Sink | 0.42 | 2.75 | 0.33 | … | 0.49 |
| Pouring Milk | 2.61 | 0.33 | 2.75 | … | 0.77 |
| … | … | … | … | … | … |
| Emptying Bottle | 0.80 | 0.49 | 0.77 | … | 2.75 |

Pairwise $S_{sim}$ Values

**B**

| Descending Order of $S_{sim}$ | Pouring Water | Washing Sink | Pouring Milk | … | Emptying Bottle |
|---|---|---|---|---|---|
| 1 | 2.75 | 2.75 | 2.75 | … | 2.75 |
| 2 | 2.75 | 2.34 | 2.75 | … | 2.62 |
| 3 | 2.71 | 1.5 | 2.75 | … | 0.94 |
| … | … | … | … | … | … |
| 15 | 0.55 | 0.31 | 0.76 | … | 0.67 |
| … | … | … | … | … | … |
| 20 | 0.21 | 0.19 | 0.17 | … | 0.22 |

Response with Highest SND is the HGT

SND row

Highest SND is the Recognition Consistency of this Video

**Fig. S1.1** Illustration of the HGT calculation for the 'pouring milk' video in Experiment 1. A) The pairwise Semantic Similarity was computed between all 20 responses for the video. B) For each response, the Semantic Similarity with the 75th percentile (15th row) semantically closest response was selected as the Semantic Neighborhood Density (SND) measure. The response with the greatest SND was defined as the HGT and the SND value was the recognition consistency for the video. See Figure 2 in the main text.

**Table S1.1**

All Action Categories Identified by Hierarchical Clustering in Experiment 1

| Action category | N | | | Mean Recognition Consistency (*SD*) | Mean $S_{sim}$ of HGTs to EK100 GTs (*SD*) |
|---|---|---|---|---|---|
| | *Easy* | *Hard* | Total | | |
| putting / placing / turning on | 19 | 39 | 58 | 0.638 (0.153) | 1.138 (0.663) |
| closing | 11 | 11 | 22 | 1.433 (0.934) | 1.189 (0.332) |
| taking | 11 | 10 | 21 | 0.648 (0.167) | 1.058 (0.501) |
| washing / drying | 10 | 11 | 21 | 1.441 (0.557) | 1.227 (0.728) |
| opening | 9 | 11 | 20 | 1.393 (0.910) | 0.986 (0.384) |
| peeling / cutting / ripping | 9 | 7 | 16 | 0.865 (0.416) | 0.996 (0.633) |
| pouring | 7 | 6 | 13 | 1.061 (0.809) | 1.178 (0.663) |
| picking / grabbing / collecting | 5 | 6 | 11 | 0.550 (0.117) | 0.734 (0.569) |
| mixing | 5 | 2 | 7 | 0.865 (0.341) | 1.243 (0.332) |
| removing | 2 | 5 | 7 | 0.495 (0.985) | 0.728 (0.472) |
| frying / microwaving | 4 | 1 | 5 | 0.719 (0.399) | 0.377 (0.181) |
| filling / emptying | 1 | 3 | 4 | 0.615 (0.144) | 0.438 (0.264) |
| getting | 4 | 0 | 4 | 0.612 (0.093) | 0.595 (0.148) |
| stirring / shaking | 1 | 3 | 4 | 1.067 (0.983) | 1.041 (0.559) |
| hanging | 2 | 1 | 3 | 1.754 (0.151) | 1.401 (0.773) |
| moving / approaching | 1 | 2 | 3 | 0.527 (0.038) | 1.352 (0.579) |
| breaking | 1 | 1 | 2 | 1.071 (0.006) | 1.254 (0.401) |
| dishing | 2 | 0 | 2 | 0.552 (0.123) | 0.376 (0.071) |
| making / preparing | 2 | 0 | 2 | 0.530 (0.167) | 0.400 (0.046) |
| packing / rearranging | 1 | 1 | 2 | 0.553 (0.054) | 0.439 (0.179) |
| plugging | 0 | 2 | 2 | 0.768 (0.316) | 1.494 (0.174) |
| turning off | 1 | 1 | 2 | 1.054 (0.207) | 1.338 (0.456) |
| wiggling / flipping | 1 | 1 | 2 | 0.536 (0.137) | 1.469 (1.812) |
| wrapping /lining | 1 | 1 | 2 | 0.531 (0.228) | 0.719 (0.758) |
| brewing | 1 | 0 | 1 | 0.546 (0.000) | 0.402 (0.000) |
| folding | 0 | 1 | 1 | 0.764 (0.000) | 0.569 (0.000) |
| lifting | 0 | 1 | 1 | 0.486 (0.000) | 1.628 (0.000) |
| using | 0 | 1 | 1 | 0.409 (0.000) | 0.411 (0.000) |

### S2. Mid-level Feature Activation Scores

Graph-Based Visual Saliency (GBVS) was used to compute the mid-level feature activation scores, as this algorithm has previously been demonstrated to accurately predict human fixation locations (Harel et al., 2006). Eye fixations can serve as proxies of feature importance for a given task, hence we assessed which components of GBVS may bear the highest importance for action recognition. GBVS was implemented in MATLAB by Harel et al. (2006, archived at: https://github.com/Pinoshino/gbvs). Feature map computation was performed at resolution level 3, using four Gabor-filter orientations (0°, 45°, 90° and 135°), four motion directions (0°, 45°, 90° and 135°), and default settings for all other feature parameters.

**Table S2.1**

GBVS Feature Channels Defined

| Feature | Definition |
|---|---|
| DKLcolor | Chromatic contrast in the Derrington-Krauskopf-Lennie color space |
| Intensity | Luminance (brightness) variations |
| Orientation | Local edge orientation energy |
| Color | Double-opponent Red–Green / Blue–Yellow contrasts |
| Flicker | Frame-to-frame luminance change (temporal contrast) |
| Contrast | Local brightness variability |
| Motion | Motion energy between consecutive frames derived from optic flow |

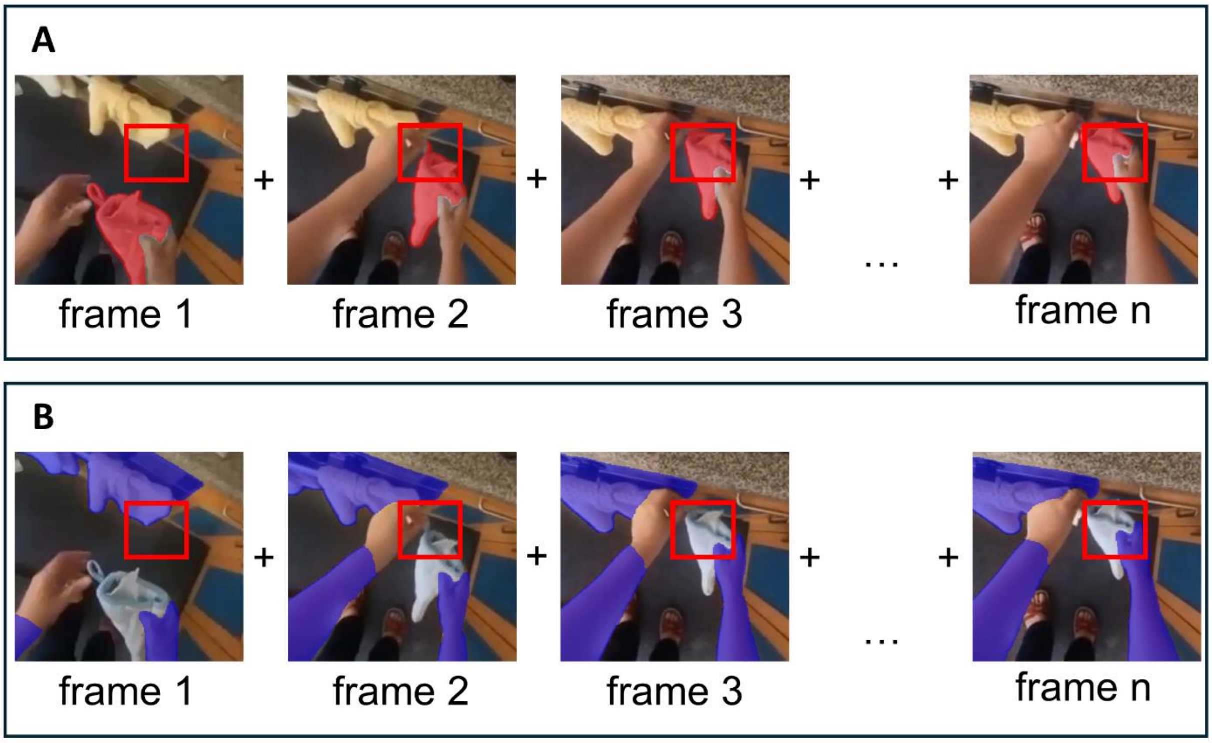


**Fig. S2.1** Examples of high-level features within a quadrant for the ‘hanging gloves’ video. A) The value for the high-level Active Object (red mask) feature was the mean proportion of pixels belonging to the segmented object contained within the quadrant (red box) relative to the full frame, averaged across frames. B) The value for high-level Contextual Objects (purple mask) feature was the mean proportion of pixels belonging to the segmented objects contained within the quadrant (red box) relative to the full frame, averaged across frames.

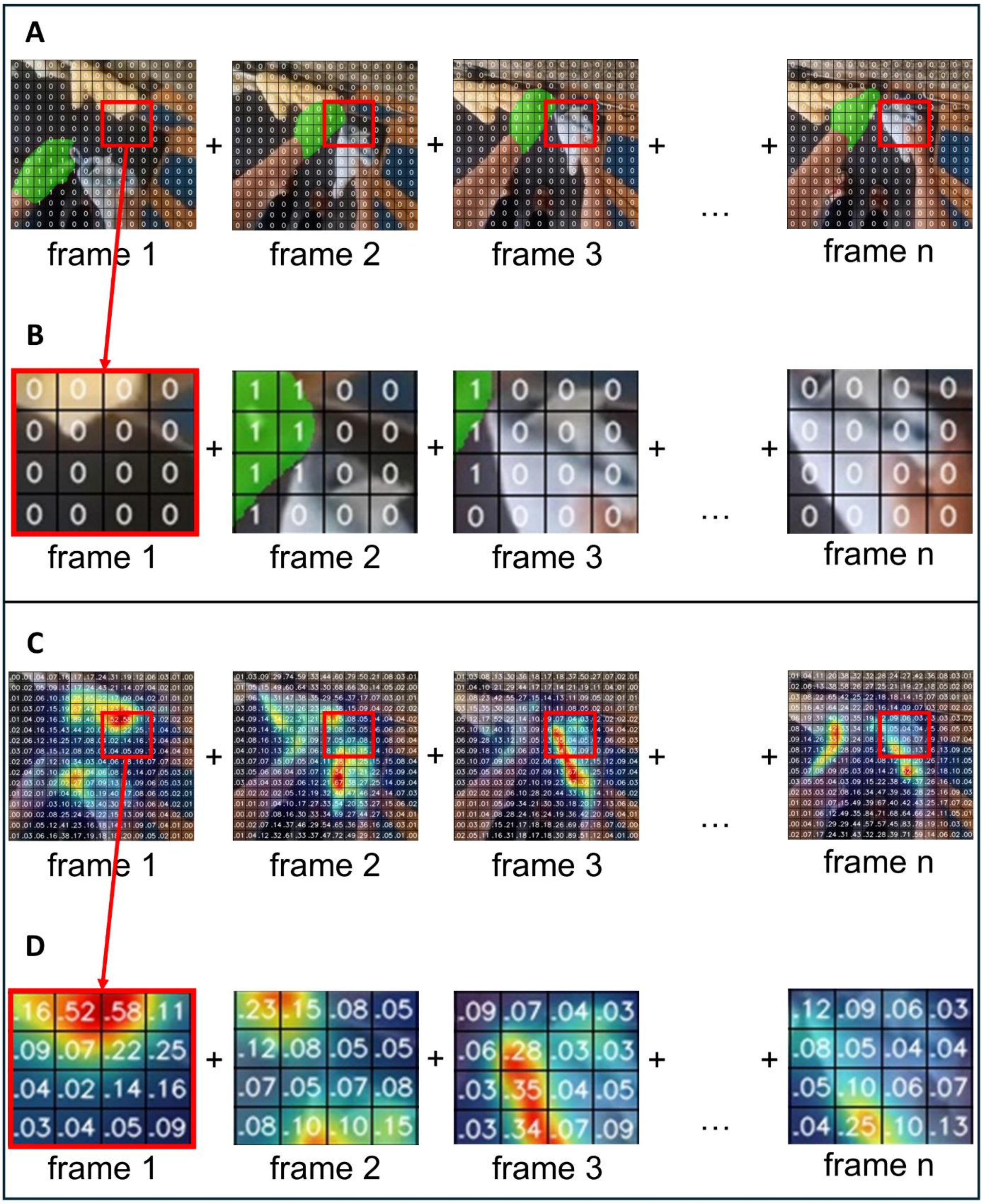


**Fig. S2.2** Detailed illustration of high-level (Active Hand) and mid-level feature (Orientation) values within a quadrant for the 'hanging gloves' video. A, B) On each frame, each pixel in the quadrant (red box; enlarged in Panel B) was determined to belong to the surface area of Active Hand (1) or not (0) based on the segmentation. C, D) On each frame, GBVS was used to compute the Orientation activation value for each pixel in the quadrant (red box; enlarged in Panel D). Activation values varied between 0 and 1 (color scale: cool = low activation, warm = high activation). The grid lines are provided to aid visualization and do not represent a single pixel. (See Fig. 8 in main text).

**S3. Quadrant Pruning**

Due to the exponential increase in number of quadrants produced through the reduction levels in Phase 1 of Experiment 2, we implemented a pruning procedure based on quadrant overlap. This helped us efficiently present only the most informative quadrants and find MIRCs more quickly.

At each data collection round, we created a selection pool of candidate quadrants. The following assumptions were applied in the following order to select quadrants from the pool for the current data collection round ($n$):

After completing data collection round $n$ - 1, we imposed overlap rules on the children generated from recognized quadrants, to decide which child quadrants would move on into the selection pool $n$ for data collection round $n$.

1. We assumed that a quadrant is unrecognizable (i.e., <50% recognition) if it is contained by >95% within a larger unrecognizable quadrant from any previous reduction levels. Therefore, we did not test it. These assumed unrecognizable quadrants did not have a valid recognition accuracy but were used in the feature learning (quadrant Type 1).
2. We merged pairs of quadrants, $Q_1$ and $Q_2$, that overlapped previously not recognized quadrant by >65% and <95% and each other by >80% and <95%. $Q_1$ and $Q_2$ were always from the same reduction level. The merged quadrant, $Q_{merged}$, was placed in selection pool $n$. If $Q_{merged}$ was unrecognizable in current data collection, $Q_1$ and $Q_2$ were assumed to be unrecognizable quadrants, with no valid accuracy. If $Q_{merged}$ was recognizable, $Q_1$ and $Q_2$ would be placed in selection pool for the next data collection round (i.e., round $n$ + 1). Only $Q_1$ and $Q_2$ were used in the feature learning process (quadrant Type 2).
3. We merged quadrants $Q_1$, $Q_2$, $Q_3$, ... $Q_n$ from single reduction level that overlapped each other by more than 95% into the merged quadrant $Q_{merged}$. One quadrant from $Q_{merged}$ was randomly selected and placed in selection pool $n$ (quadrant Type 3). The other individual quadrants were assumed to have the same recognition accuracy as $\mathrm{Q}_{\mathrm{merged}}$ (quadrant Type 4). The individual quadrants were used for feature learning.
4. All quadrants that did not meet criteria 1-3 were placed in selection pool $n$.

After generating the selection pool $n$ for the current data collection round, we applied the following rules to select final quadrants to be tested in data collection round $n$:

1. All merged quadrants were tested.
2. All (non-merged) quadrants that overlapped a larger unrecognizable quadrant from any previous levels by 65% or more were tested (quadrant Type 0).
3. We ranked the remaining quadrants by their cumulative area overlap with other quadrants in the selection pool $n$. Cumulative area overlap of quadrant $Q_1$ was calculated by summing the proportion of $Q_1$'s area covered by quadrants $Q_2$, $Q_3$, $Q_4$, ... $Q_n$ of that video. For each video we then selected as many highest scoring quadrants as possible to end up with equal number of quadrants per video in data collection round $n$. This allowed us to maximize the number of data points per participant and maximize the amount of information available for the pruning procedure of future data collection rounds (quadrant Type 0).

4. All unselected quadrants were shifted into selection pool $n + 1$.

The data collection rounds tested recognition of 4521 merged and regular quadrants. The pruning procedure was applied for data collection rounds D, E and F. Data collection rounds A, B and C involved all available quadrants, while for data collection round G we imposed overlap rules and from the selection pool hand-picked quadrants, whose unrecognizability would confirm the MIRC status of quadrants for videos that lacked a MIRC (N = 2) or had a low number of MIRCs (N = 3). These MIRC candidates were quadrants which already had 3 children confirmed as being unrecognizable, lacking only 1 more. Before applying the pruning procedure, selection pool 8 terminated at 2826 individual quadrants.

**Table S3.1**

Characteristics of Quadrant Types

| Quadrant Type | N quadrants | Has valid recognition accuracy | Used for feature learning |
|---|---|---|---|
| 0 | 3427 | YES | YES |
| 1 | 2235 | NO | YES |
| 2 | 643 | NO | YES |
| 3 | 565 | YES | YES |
| 4 | 734 | YES | YES |

*Note.* Quadrant Types 1 and 2 do not have a valid recognition accuracy because they were assumed to be unrecognizable based on their overlap with other unrecognizable quadrants.

**Table S3.2**

Characteristics of Data Collection Rounds in Phase 1 of Experiment 2

| Data collection round | Participants | MIRCs | non-MIRCs | |
|---|---|---|---|---|
| | | | Recognizable | Unrecognizable |
| A | 80 | 7 | 123 | 14 |
| B | 320 | 23 | 317 | 180 |
| C | 1280 | 204 | 627 | 529 |
| D | 820 | 120 | 269 | 509 |
| E | 680 | 93 | 221 | 301 |
| F | 580 | 27 | 198 | 220 |
| G | 40 | 0 | 1 | 9 |

*Note.* Data collection rounds D-G included quadrants from various Reduction Levels.

### S4. MIRCs from Experiment 2

**Table S4.1**

Distribution of MIRCs Across *Easy* Videos

| Video ID | HGT | MIRCs |
|---|---|---|
| 3159 | Turning off light | 28 |
| 4376 | Closing cabinet | 10 |
| 9152 | Hanging gloves | 14 |
| 10766 | Pouring milk | 14 |
| 20586 | Putting water | 18 |
| 27161 | Filling water pot | 14 |
| 28393 | Closing dishwasher | 2 |
| 29776 | Opening oil bottle | 27 |
| 36270 | Taking out plate | 16 |
| 37869 | Putting away dishcloth | 7 |
| 41842 | Washing cutting board | 4 |
| 48868 | Taking out plate | 24 |
| 50987 | Opening cabinet | 19 |
| 51854 | Washing plate | 16 |
| 57408 | Opening washing machine | 7 |
| 59895 | Cutting vegetable | 22 |
| 63504 | Taking tissue paper | 20 |
| 65460 | Putting away vacuum | 11 |

*Note*. ‘Video ID’ refers to ID in the Epic-Kitchens-100 dataset (Damen et al., 2022).

**Table S4.2**

Distribution of MIRCs Across *Hard* Videos

| Video ID | HGT | MIRCs |
|---|---|---|
| 10788 | Opening milk | 13 |
| 15469 | Putting broom away | 10 |
| 16092 | Opening food packet | 19 |
| 23533 | Turning off switch | 3 |
| 23590 | Pouring something | 16 |
| 23843 | Open oil container | 9 |
| 26714 | Taking scissors | 6 |
| 30735 | Scrubbing pot | 9 |
| 32600 | Putting into fridge | 5 |
| 33974 | Taking coffee mug | 9 |
| 34562 | Taking out container | 21 |
| 34567 | Closing cabinet | 5 |
| 35778 | Puring liquid | 23 |
| 44943 | Peeling of carrot | 12 |
| 46211 | Hang cloth | 18 |
| 55836 | Cleaning knife | 5 |
| 58280 | Putting spoons away | 9 |
| 62570 | Closing container | 9 |

*Note*. ‘Video ID’ refers to ID in the Epic-Kitchens-100 dataset (Damen et al., 2022).

### S5. Spatiotemporal MIRCs vs. Spatial MIRCs

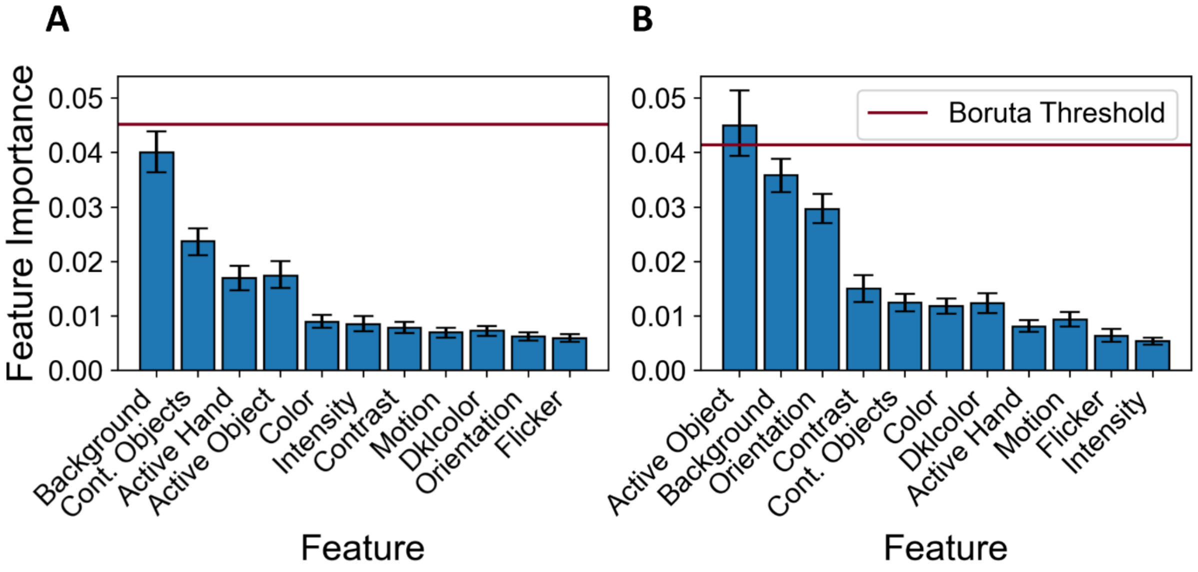


**Fig. S5.1** Feature importance for the non-significant binary classification in Experiment 2. A) Mean feature importance for the classification between spatial vs spatiotemporal MIRCs from *Easy* videos. B) Mean feature importance for the classification between spatial and spatiotemporal MIRCs from *Hard* videos. Mean feature importance greater than the Boruta Threshold (purple) were significantly informative for the classifier. Features are ordered by their SHAP summaries in Figure S6.1. Error bars reflect 95% confidence intervals.

**Table S5.1**

Mean Feature Values for *Easy* and *Hard* Spatial and Spatiotemporal MIRCs

| Feature | *Easy* | | *Hard* | |
|---|---|---|---|---|
| | Spatial | Spatiotemporal | Spatial | Spatiotemporal |
| Active Object | 0.160 (0.145) | 0.013 (0.012) | 0.314 (0.249) | 0.196 (0.169) |
| Background | 0.029 (0.036) | 0.051 (0.065)* | 0.030 (0.036) | 0.041 (0.048) |
| Contextual Objects | 0.077 (0.063) | 0.067 (0.065) | 0.117 (0.112) | 0.111 (0.114) |
| Active Hand | 0.094 (0.135) | 0.097 (0.014) | 0.156 (0.199) | 0.146 (0.199) |
| Color | 0.076 (0.054) | 0.084 (0.060) | 0.084 (0.042) | 0.086 (0.070) |
| Intensity | 0.083 (0.043) | 0.093 (0.060) | 0.085 (0.037) | 0.088 (0.060) |
| Contrast | 0.077 (0.046) | 0.093 (0.070) | 0.075 (0.037) | 0.089 (0.077) |
| Motion | 0.086 (0.053) | 0.087 (0.057) | 0.105 (0.056) | 0.104 (0.075) |
| DKLcolor | 0.081 (0.049) | 0.084 (0.061) | 0.100 (0.050) | 0.095 (0.060) |
| Orientation | 0.100 (0.061) | 0.113 (0.072) | 0.126 (0.053) | 0.123 (0.081) |
| Flicker | 0.079 (0.044) | 0.084 (0.055) | 0.099 (0.052) | 0.098 (0.069) |

*Note. SD* is in parenthesis. There were no significantly informative features for the binary classification spatial vs spatiotemporal MIRCs for both datasets, as classifier did not reach above-chance performance. Asterisk '*' indicates a significant difference in mean feature value between spatial and spatiotemporal MIRCs within the given dataset.

**6. SHAP Summary Plots**

The SHAP summary plots are based on single classification run without shadow features (Lundberg & Lee, 2018). A SHAP summary plot ranks features in descending order of importance (mean absolute SHAP value averaged across stimuli). Each point represents SHAP value of that feature for a single video. Positive SHAP values indicate influence towards a class-1 decision (e.g., ‘MIRC’), negative SHAP values represent influence towards a class-0 decision (e.g., ‘Unrecognized Quadrant’). The color represents the feature value within a quadrant relative to the full frame, with cool colors representing low values of that value (e.g., low proportion of the active object’s surface area visible in a quadrant) and warm colors representing high values of that feature (e.g., high proportion of the active object’s surface area visible in a quadrant). SHAP summary plots are presented in Figures S6.1A-E.

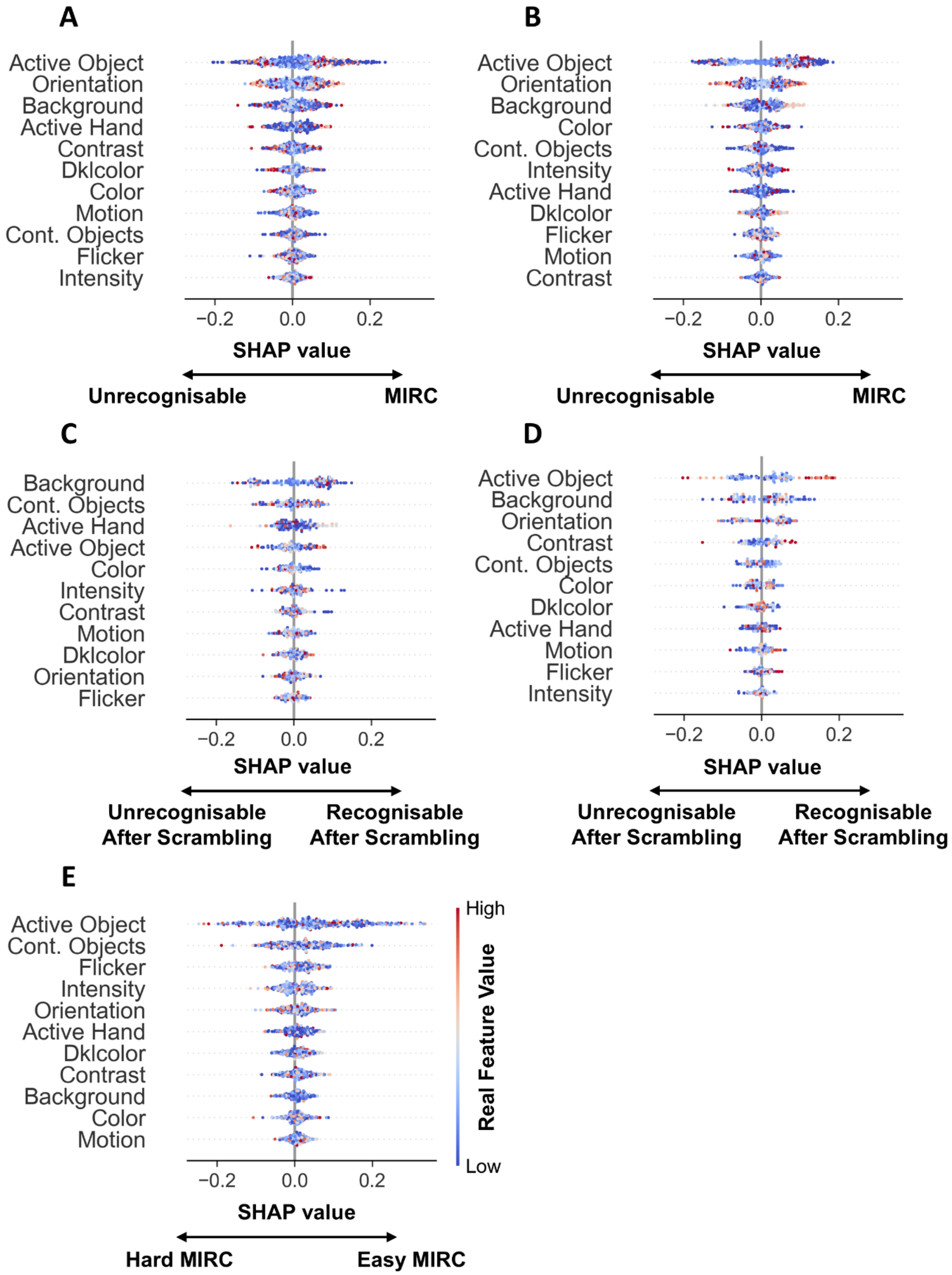


**Fig. S6.1** SHAP summary plots for all binary classifiers in Experiment 2. A) SHAP summary for classification between MIRCs and unrecognizable quadrants from *Easy* videos. B) SHAP summary for classification between MIRCs and Unrecognizable Quadrants from *Hard* videos. C) SHAP summary for classification between Spatial and Spatiotemporal MIRCs from *Easy* videos. D) SHAP summary for classification between Spatial and Spatiotemporal MIRCs from *Hard* videos. E) SHAP summary for classification between *Easy* and *Hard* MIRCs.
